# Inferring active and passive mechanical drivers of epithelial convergent extension

**DOI:** 10.1101/2025.01.28.635314

**Authors:** Sommer Anjum, Deepthi Vijayraghavan, Rodrigo Fernandez-Gonzalez, Ann Sutherland, Lance Davidson

## Abstract

What can we learn about the mechanical processes that shape tissues by simply watching? Several schemes suggest that static cell morphology or junctional connectivity can reveal where chains of cells transmit force or where force asymmetries drive cellular rearrangements. We hypothesize that dynamic cell shape changes from time lapse sequences can be used to distinguish specific mechanisms of tissue morphogenesis. Convergent extension (CE) is a crucial developmental motif wherein a planar tissue narrows in one direction and lengthens in the other. It is tempting to assume that forces driving CE reside within cells of the deforming tissue, as CE may reflect a variety of active processes or passive responses to forces generated by adjacent tissues. In this work, we first construct a simple model of epithelial cells capable of passive CE in response to external forces. We adapt this framework to simulate CE from active anisotropic processes in three different modes: crawling, contraction, and capture.

We develop an image analysis pipeline for analysis of morphogenetic changes in both live cells and simulated cells using a panel of mechanical and statistical approaches. Our results allow us to identify how each simulated mechanism uniquely contributes to tissue morphology and provide insight into how force transmission is coordinated. We construct a MEchanism Index (MEI) to quantify how similar live cells are to simulated passive and active cells undergoing CE. Applying these analyses to live cell data of *Xenopus* neural CE reveals features of both passive motion and active forces. Furthermore, we find spatial variation across the neural plate. We compare the inferred mechanisms in the frog midline to tissues undergoing CE in both the mouse and fly. We find that distinct active modes may have different prevalences depending on the model system. Our modeling framework allows us to gain insight from tissue timelapse images and assess the relative contribution of specific cellular mechanisms to observed tissue phenotypes. This approach can be used to guide further experimental inquiry into how mechanics influences the shaping of tissues and organs during development.

## INTRODUCTION

Embryonic epithelial tissues undergo shape changes during development to form tissues and organs. Time lapse sequences of tissue morphogenesis may encode the cellular and physical processes that shape them. Force inference methods based on mechanical models of force balance propose that cell shapes and morphometry such as the lengths and angles of cellular junctions reveal patterns of cell-generated force (Brodland et al. 2014; Ishihara and Sugimura 2012; Merkel et al. 2016; Guirao et al. 2015; Huebner et al. 2021; Claussen, Brauns, and Shraiman 2024). Force inference models can serve as hypotheses of force patterns in tissues, but they are not designed to offer insights into specific mechanisms behind the collective cell movements driving tissue morphogenesis. The goal of this work is to develop techniques that can be applied to time lapse sequences of epithelia such that we can infer cell-level mechanisms driving tissue shape change.

Epithelia come in many forms, but the simple epithelia we consider here consist of a monolayer of cells physically connected into a flat sheet across their apical surfaces (Eurell and Frappier 2013; Banks 1993; Ganz 2002; Davies and Garrod 1997). These multicellular sheets serve as a barrier to the environment that persists during the dynamic processes of morphogenesis including cell proliferation, apoptosis, intercalation, and cell rearrangement (Powell 1981; Eisenhoffer and Rosenblatt 2013; Lemke and Nelson 2021). Furthermore, the epithelium’s barrier function needs to be robust to environmental perturbations such as pressure, tension, shear, and harsh pH levels (Gibson et al. 1996; Macara et al. 2014; McNeil 1993). These functional requirements are especially critical when epithelia undergo rapid or large deformations of development to form tissues and organs.

A key example of an epithelial morphogenetic process that fulfills these functional requirements is convergent extension (CE). During CE, a tissue narrows along one axis and extends in the orthogonal axis within the plane of the epithelial sheet (Walck-Shannon and Hardin 2013; Keller et al. 2000; Warga and Kimmel 1990; Fernandez-Gonzalez et al. 2009). CE is a conserved process in the morphogenesis of elongated structures. Examples include CE during *Drosophila* germ band elongation (Rauzi, Lenne, and Lecuit 2010; Bertet, Sulak, and Lecuit 2004; Irvine and Wieschaus 1994), neurulation in vertebrates (Davidson and Keller 1999; Butler and Wallingford 2018; Keller et al. 2000; Harris and Juriloff 2010; Vijayraghavan and Davidson 2017; Williams et al. 2014), and elongation of the kidney during human development (Torban and Sokol 2021; Lienkamp et al. 2012). CE shapes the early neural tube, the precursor of the central nervous system. Defects in neural CE can result in developmental disorders with high prevalence, such as spina bifida and anencephaly which have an incidence of 0.5 per 1000 live births in the US every year (Kancherla, Redpath, and Oakley 2019; Vijayraghavan and Davidson 2017). It is critical to understand how cellular movements are coordinated during CE so that we can understand where epithelial mechanical errors emerge.

To shape tissues and organs during development, cellular force generation must be coordinated both spatially and temporally. Forces that shape embryonic epithelia are largely the product of the intracellular dynamic cytoskeleton, namely contraction of the actomyosin meshwork in the apices of polarized epithelia (Pannekoek, de Rooij, and Gloerich 2019; Coravos, Mason, and Martin 2017). Force generated by contraction in one cell can be propagated to adjacent cells via the mechanical coupling of the adherens junction, which connects to the cytoskeleton (Harris, Daeden, and Charras 2014; Charras and Yap 2018; Ng et al. 2014). Molecular players are involved in the feedback between physical and biochemical processes. Biochemical drivers include planar cell polarity (PCP) proteins that establish the directionality of force generation machinery and components of the adherens junctions that remodel in response to force (Wang et al. 2019). Mutations in cytoskeletal regulatory proteins are associated with neural tube defects in mammals and humans (Harris and Juriloff 2010; Dubielecka et al. 2011; Haigo et al. 2003). The mechanical roles, e.g. in their effects on mechanical properties, force-generation, or force-transmission of these defects have been challenging to isolate experimentally. To overcome these challenges, we adopt a data-driven computational modeling approach. We hypothesize that representing proposed mechanisms of tissue shaping at the cellular level in a computational model will result in variations in tissue morphology. Quantifying these variations will allow us to identify potential cellular mechanisms from timelapse data. Using this system, we hope to infer mechanisms of CE and prioritize hypotheses to test *in vivo*.

To simulate epithelial CE, force generation must be explicitly represented and then transmitted between cells to change the shape of the tissue. Narrowing one tissue axis and extending the other during CE requires an imbalance of forces, or asymmetry in tissue mechanics (Simões, Mainieri, and Zallen 2014; Keller and Sutherland 2020; Wang et al. 2020a; Ioratim-Uba, Liverpool, and Henkes 2023). Enhanced contractility along one tissue axis, differential mechanical resistance, and asymmetric activation of adherens junctions have all been suggested to introduce this anisotropy (Shindo 2018; Solnica-Krezel and Cooper 2002). Four common computational frameworks have been used to simulate the epithelial surface: Vertex models (Staple et al. 2010; Fletcher et al. 2014; Hardin and Weliky 2019; Barton et al. 2017), Cellular Potts models (Graner and Glazier 1992; Wolff, Davidson, and Merks 2019), Finite Element models (Brodland and Veldhuis 2012), and Active Particle models (Dalle Nogare and Chitnis 2020; Sulsky, Childress, and Percus 1984a; Bi et al. 2016). Other models exist outside these frameworks, and we direct the reader to the references for more examples (Moure and Gomez 2021; Meineke, Potten, and Loeffler 2001; Amonlirdviman et al. 2005; Vangheel, Ongenae, and Smeets 2022). Cell-generated forces can be represented in vertex models by variations on a line tension term representing actomyosin contractility (Staddon et al. 2019).

Such contractions can contribute to ratcheting of the apical surface by creating a line tension term that is time-dependent for horizontal junctions. In another vertex model, nonlinear feedback of tension increases to continuously deform the tissue (Claussen, Brauns, and Shraiman 2024). Vertex models can also be used to encode junctions of neighboring cells with both a pushing force and a contraction force (Weng, Huebner, and Wallingford 2022; Cavanaugh et al. 2020).

Similarly, in a Cellular Potts framework, CE can be achieved by endowing each cell with links representing polarized filopodial protrusions that exert forces between the centers of connected cells (Belmonte, Swat, and Glazier 2016). Another approach assigns force-generating filaments to lattice elements (Rens and Edelstein-Keshet 2019). Alternatively, a Cellular Potts model was employed to demonstrate that anisotropic differential adhesion alone was sufficient to drive CE (Zajac, Jones, and Glazier 2003). To mimic discrete force-generation in a finite element scheme, a model of neurulation achieves force asymmetry via a lamellipodial stress between cells (Chen and Brodland 2000). The same effect in a finite element model can be achieved with force-generating rod elements (Brodland, Viens, and Veldhuis 2007). Active particle models with cells represented by a single node are not typically used to simulate CE. Nonetheless, forces can be defined asymmetrically on a cell-to-cell basis, with cells being self-propelled (Yang et al. 2017). The resulting network of cell-centroid nodes is then tessellated into an epithelial sheet.

Thus, asymmetric cell-generated forces can be implemented in active particle, vertex, and finite element models where each are capable of simulating CE. To ascribe forces on a cell-to-cell basis with limited assumptions on cell adhesion and remodeling, we chose to use an active particle modeling framework (Manning and Collins 2015; Anjum et al.).

The aim of this study is to develop passive and active modes in a common CE simulation framework so that we can identify distinct features of cell- and tissue-dynamics that are correlated to the driving mode. We are further guided by live imaging and biophysical studies of CE during *Xenopus* neurulation (Davidson and Keller 1999; Davidson et al. 2006; Elul and Keller 2000; Keller et al. 2000) and during germband elongation in *Drosophila* (Bertet, Sulak, and Lecuit 2004; Sun et al. 2017; Wang et al. 2020b; Irvine and Wieschaus 1994; Zallen and Wieschaus 2004; Blankenship et al. 2006) The simplest hypothesis is that CE in these cases may be a response to forces from outside, or external to, the epithelium (Sethi, Cram, and Zaidel-Bar 2017; Collinet et al. 2015; Lye et al. 2015). We refer to this as the tissue being shaped by “passive forces”. Conversely, epithelial cells can also generate their own force to drive CE. We refer to this as the tissue being shaped by “active forces”. We simulate three different mechanisms of active force generation that are proposed to be involved in CE: contraction of ML junctions, cells crawling toward the midline, and midline capture (inspired by boundary capture in the mesoderm (Reintsch et al. 2005)). Quantification of the differences in cellular and tissue shape change across passive and active modes will allow for “mechanism inference”. By using a palette of metrics inspired by solid mechanics, soft matter physics, and shape-based morphometrics, we define criteria to guide mechanism inference of each mode of CE. We apply the same analyses to live cell data from CE in frog, fly, and mouse so that we might predict force profiles present *in vivo*. Our mechanism inference approach provides a framework to assess cellular mechanisms driving the shaping of tissues and organs and to prioritize hypotheses for further experimental inquiry into the mechanics of morphogenesis.

## RESULTS

### Convergent extension of the neural epithelium in *Xenopus*

Our computational models are inspired by CE during neurulation in *Xenopus laevis* embryos. In *Xenopus*, the most pronounced phase of CE occurs in the flat neural plate in the trunk from the conclusion of gastrulation (NF 13) through mid-neurula stages (NF 15) (Fig. 1A-B, Supplemental Video 1; brightfield timelapse). During CE, cells in the neural epithelium undergo neighbor exchanges as the tissue elongates. Cells that are neighbors lose the junction connecting them in the anteroposterior (AP) direction and acquire new neighbors through the formation of a new orthogonal junction in the mediolateral (ML) direction (Fig. 1C, Supplemental Video 2; confocal timelapse). Such neighbor exchanges are referred to as a “T1 transition” (Fig. 1C), after studies of soap foams and other granular soft matter (Jain, Voigt, and Angheluta 2024; Spencer, Jabeen, and Lubensky 2017; Weaire, Barry, and Hutzler 2010). Mediolaterally directed neighbor exchanges are common as strain rates reach 15% strain per hour as the neural plate elongates anteroposteriorly.

**Figure 1.**
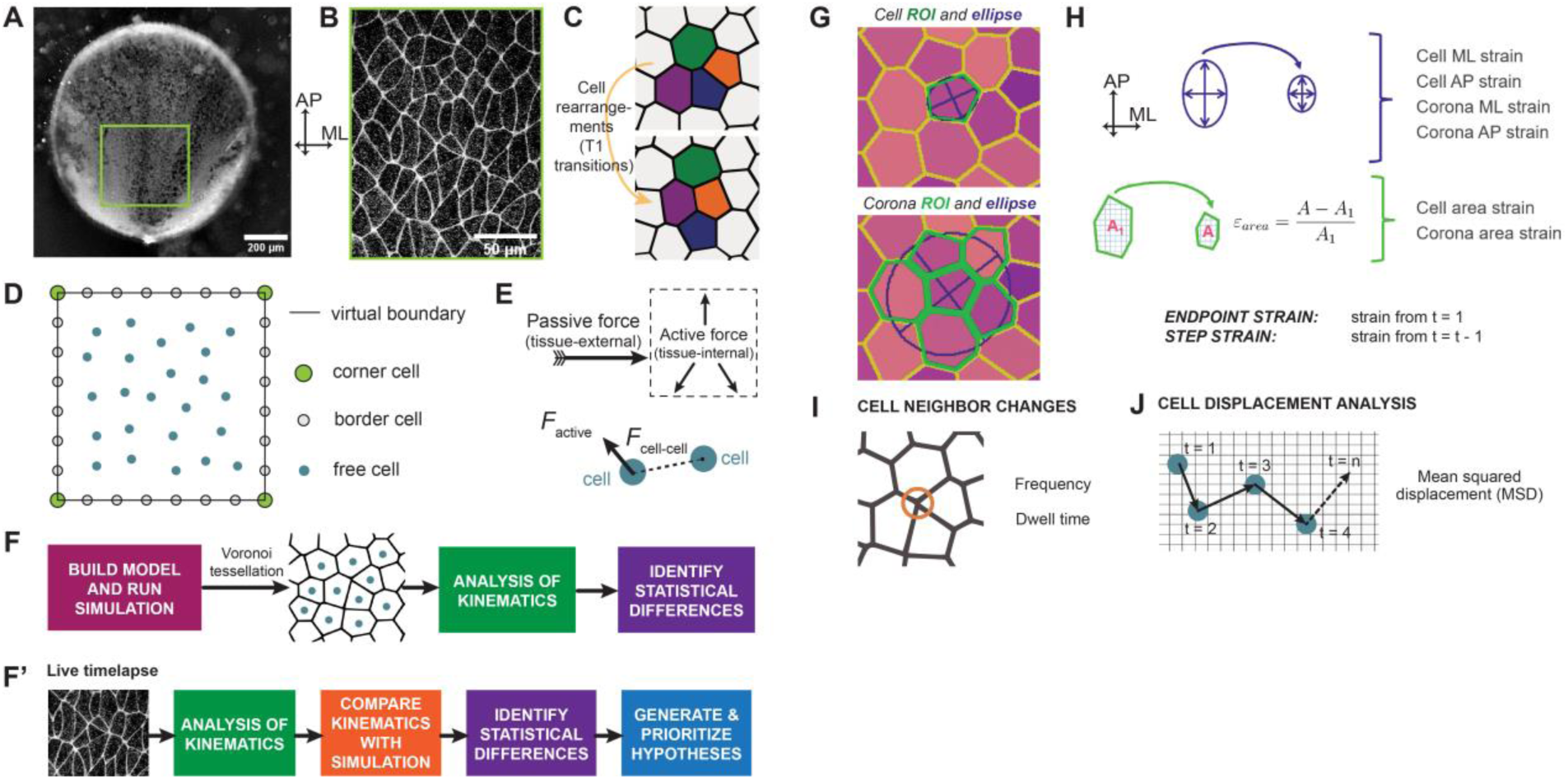
Quantitative dynamics of convergent extension (CE) in simulations and live cell timelapses. (A) Representative frame from a *Xenopus* whole-embryo brightfield timelapse with the neural plate, posterior of the hindbrain, indicated with a green box (see also Supplementary Video 1). (B) Representative frame from a confocal timelapse from a posterior region of the *Xenopus* neural plate labeled with CAAX-GFP (see also Supplementary Video 2). (C) Pictorial representation of cell neighbor exchanges that are characteristic of CE. (D) Simulation schematic of the centers of different cell types in the model. (E) Definition of passive and active forces and diagram of how forces act between two free cell centroids. (F) Steps to perform simulation and extract data to identify signatures. (F’) How steps from E are applied to live cell data to extract signatures. (G) Designation of cell and corona regions of interest (ROIs) and how they are used to calculate different types of strains. (H) Description of how cell and corona shapes are used to calculate strains. (I) Cell neighbor change analysis for frequency and dwell time of cells in transition states of higher order/T1 junctions. (J) Schematic of concepts behind mean square displacement (MSD) calculation.

### A simple model of an epithelium in which passive and active forces can be represented independently

To simulate epithelial CE, we adopt a cell centroid-, or node-based model with free cells, border cells, and corner cells (Fig. 1D). Each type of cell has its own movement rules. Free cells are represented by interior nodes that move in both axes within the 2D domain. The 2D domain itself is bounded by fixed corner cells and border cells that move freely along the border of the 2D domain. Passive forces external to the tissue can be applied as displacements to the nodes that define the 2D domain. Active tissue-internal forces are simulated through interactions between free cells (Fig 1E). Free cells are mutually repulsive if they are sufficiently close, exerting a force F_cell-cell_ that maintains node spacing. As the borders move, free and border cells rearrange in response to the sum of F_cell-cell_ from their neighbors. Passive and active forces can be represented independently in the equation of motion for each node:

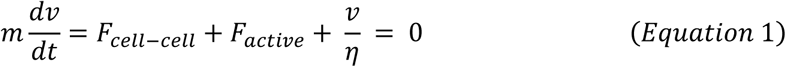

where cell-cell (F_cell-cell_), active intrinsic (F_active_) forces, and drag forces (velocity *v*, scaled by the viscosity η) are balanced in the quasi-static case of the embryo where cell motion is fully damped.

Cell-cell forces are represented by a repulsive potential from each cell center that is a function of distance, *dist*. This is separated into x and y components by calculating *dx* and *dy*, the x and y distances between cells *i* and *j*, respectively. Cell-cell forces act within an interaction distance, *rest*, of cell *i,* outside which the force is zero. Repulsion forces are defined in the x and y directions as:

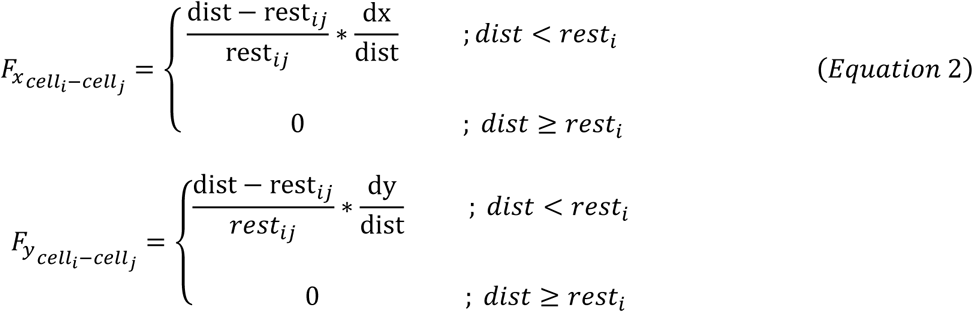

To mimic the variation in cell sizes that are observed *in vivo*, we introduce mechanical heterogeneity in the epithelium by assigning each node a repulsive potential that is randomly selected from a uniformly distributed set of potentials/rest lengths (rest_i,j,k…_). *Diameter* is used to set an average cell size, and *dist* is the distance between centroids. These quantities are used to calculate the resting distance between cells to calculate the forces in Equation 2.

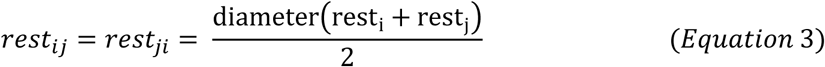

The x and y cell-cell repulsion forces are summed for all cells *j* though *n* within the interaction distance, *rest*, of cell *i*,

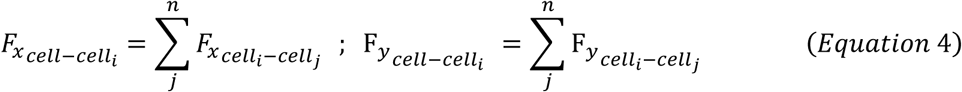

The total force in each direction is the sum of cell-cell and active forces on each node:

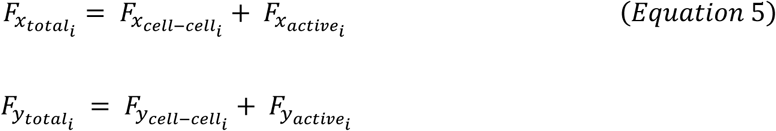

The simulation advances in time after forces on all nodes *i* are calculated, and cell positions are updated using forward Euler methods:

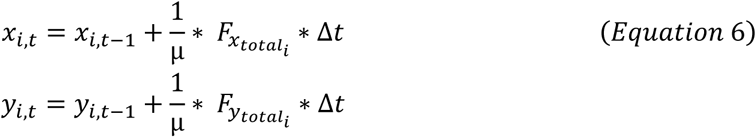

Subsequent motions are analyzed using an analysis pipeline (Fig. 1F) that can be applied to both live cell data and simulated classes of CE (Fig. 1F’). The first step of the pipeline, whether it be for live cells or simulated cells, is to perform a segmentation. For live cells, we segment cell boundaries on a fluorescent protein label of the plasma membrane (mem-RFP) (Mashburn et al. 2012); for simulations, cell boundaries are rendered by Voronoi tessellation of the centroids into a representative cell sheet (Bi et al. 2016; Sulsky, Childress, and Percus 1984b; Meineke, Potten, and Loeffler 2001). Once segmented, we identify cell shapes, or regions-of-interest (ROIs) of individual cells using open-source image analysis software (Schindelin et al. 2012). Using the trajectories of cell ROIs over time, we carry out a first “level” of analysis to quantify cell and tissue movement and dynamic shape change.

### First level analysis: tracking cell- and tissue-deformation

From changes in cell and tissue shape, we calculate strain rates throughout the time-lapse sequence. We designate a one-level “corona”, or the region containing a cell and its immediate neighbors, to track each cell’s local mechanical environment (Blanchard et al. 2009) (Fig. 1G). For all timesteps, the kinematics of the boundary containing the same set of neighbor cells are analyzed, regardless of whether neighbor changes have occurred. At each timestep, we fit an ellipse to each cell to calculate its strain, and another ellipse is fitted to its corona to calculate its strain. Mediolateral (ML) and anteroposterior (AP) strains, align to the x and y axes of the image frame, respectively, are defined in the Methods section. The area within the cell and corona ROIs are used to calculate area strain (Fig. 1H). In total, we calculate three strain values for each cell and its corona, for a total of six strain values, denoted with *B*: Cell ML, Cell AP, Corona ML, Corona AP, Cell area, and Corona area strains. We define two types of strain using these quantities:

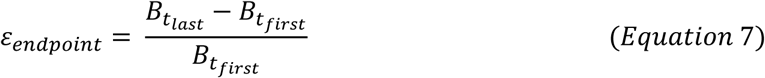

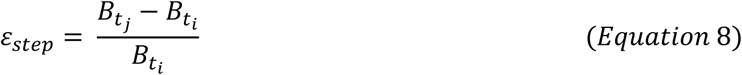

Endpoint strains allow us to measure long-term strain rates by comparing shape changes from the first (t_first_) to the last (t_last_) timestep, while step strains represent short-term shape changes between timesteps *i* and *j*. These strains can be represented as strain rates (ε̇) by dividing by the time interval over which the strain occurs; however, we refer to these simply as strains, as the time step is constant for all endpoint strains (the full time lapse) and step strains (one time step).

From segmented timelapses, we also track the formation and duration of higher-order junctions (greater than three cells meeting at a vertex). Higher-order junctions represent a transitional state of neighbor exchanges characteristic of remodeling epithelial sheets including those undergoing CE (Fig. 1I). Temporal tracking of individual junctions reveals the frequency of each cell’s participation in T1 transitions in addition to how many timesteps a cell spends in a high order junction. Tracked cell centroids also allow the calculation of mean squared displacement (MSD) in both the ML and AP axes (Fig. 1J).

### Second-level analysis: leading-lagging analysis

Step strain provides insight into how cell and tissue shape changes are temporally related. As convergent extension proceeds, there are three possibilities for cell and tissue shape change (Fig. 2A): either (i) the cell strains first, (ii) the corona strains first, or (iii) the cell and corona strain together. We can quantify which one of these possibilities are more prevalent. We compare positively phase shifted (+1 to +5 timesteps), negatively shifted (-1 to -5 timesteps), or unshifted correlations between corona and cell strain. For each phase shift, we calculate the *lead* metric, which is calculated differently based on whether the cell and corona strains have the same sign:

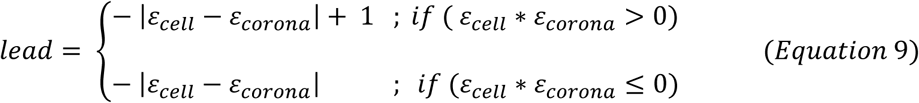

**Figure 2.**
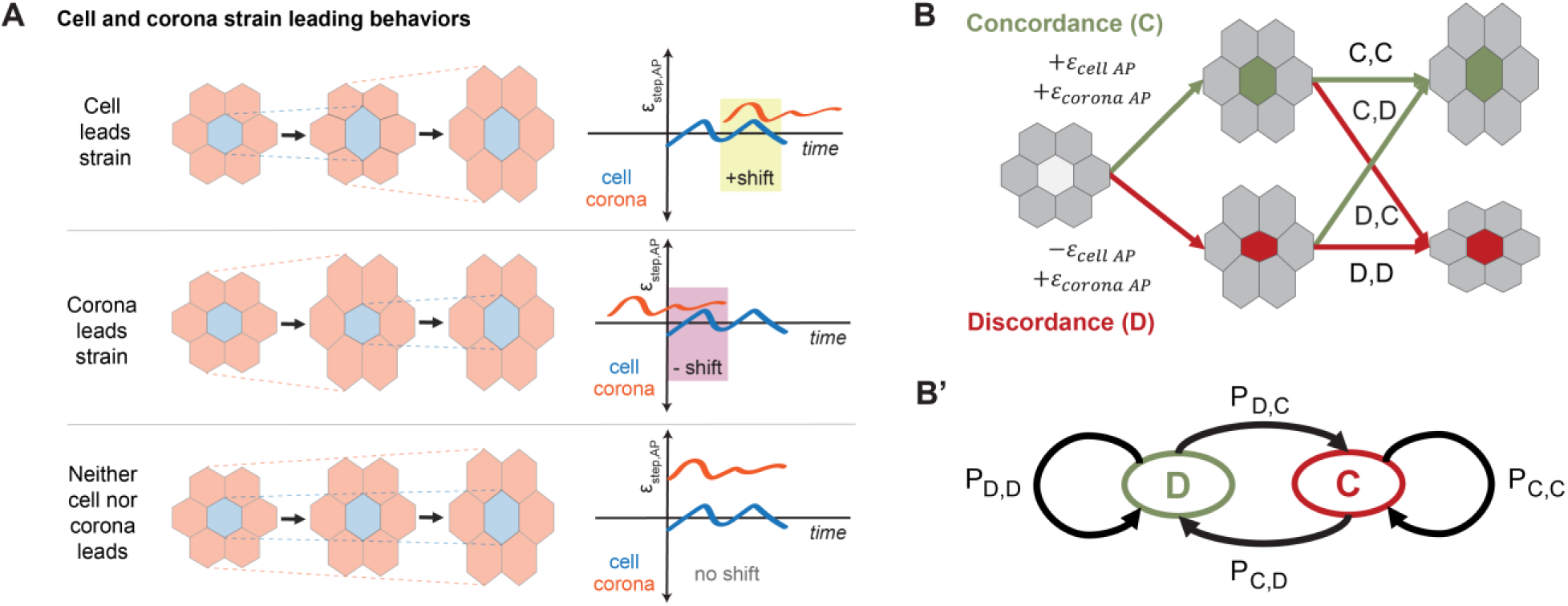
Analyses of the agreement in cell and corona strain timing. (A) There are three ways to achieve a strain configuration of the cell and corona: the cell straining first, the corona straining first, and the cell and corona straining together. Each case is shown with a pictorial representation, and sample trajectories of cell and corona strain rates for positive, negative, and zero shift are illustrated with plots. (B) Pictorial representation of how cell and corona strain rates may have different signs. (B’) How these concordant and discordant states are described with a two-state Markov model with transition probabilities between states.

The values of *lead* are averaged for all five positive shifts and all five negative shifts for comparison. Higher *lead* values indicate more correlated strains, while more negative *lead* values indicate the opposite. If the lead metric is higher for a positive shift, this indicates that the cell strains precede corona strain. Conversely, if the lead metric is higher for a negative shift, this indicates that the corona strains precede cell strain. Comparing the lead metric for different shifts of ML, AP, and area strain can inform us of how strain is being transmitted in the epithelial sheet in each mode of CE that is simulated, and we can compare this to what we observe in the live-cell data. This allows us to directly quantify local and global force transmission as a function of how it is generated.

### Second-level analysis: two-state Markov model

The lead metric begins our inquiry into spatial and temporal force transmission across cell and tissue scales. To further assess the mechanical coordination of deformation between cells and their surrounding tissue in time, we introduce *concordance* as the cell and corona retaining the same sign of strain (both positive or negative) and *discordance* as the cell and corona displaying different signs of strain (one positive, one negative) over a timestep (Fig. 2B).

Furthermore, cells may change states of concordance and discordance over time. To summarize these state changes, we calculate concordance transitions in a two-state Markov model (Fig. 2B’). Concordance probabilities using the step strain data quantify the probability of cells undergoing four possible transitions between adjacent time steps: (i) staying in concordance, (ii) staying in discordance, (iii) moving from concordance to discordance, or (iv) moving from discordance to concordance. Differences in Markov model transition probabilities can reveal differences in strain coordination for different hypothesized CE mechanisms, shedding light on the operation of these mechanisms in live cell data.

### Initial conditions: Matching simulated epithelial variation and network morphologies to live epithelial tissues

During morphogenesis *in vivo* epithelial cell morphology varies both locally and across tissues. To represent this variability within our model epithelia we first establish a set of parameters to use as a “base case”. Live cells exhibit variation in cell area and sidedness (Supplementary Figs. S1A) that have been noted previously both in live epithelia and simulated epithelial systems (Gibson et al. 2006; Staple et al. 2010). Distributions of area and sidedness were compared between simulations and experimental observations (Supplementary Fig. S1B) as parameters including cell packing density, speed of border movement, and viscosity are varied (Supplementary Table 1). From these parameter sweeps, we identified a base case set of parameters that match *in vivo* area and sidedness distributions (Supplementary Table 2).

### Convergent extension by externally applied force

How epithelia remodel under externally applied force is not well understood. In order to apply external forces to our model, we simulate CE by moving the left and right borders inward along the x-direction and moving the top and bottom borders apart in the y-direction such that area is conserved (Supplementary Fig. S2, Supplemental Video 3). The length and width of the boundaries change based on a rate (pixels per timestep of the simulation):

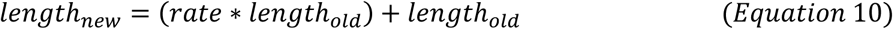

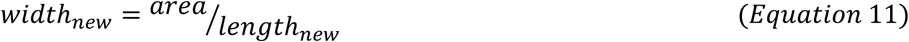

The simulation is stopped once strain in the AP direction reaches 15% as determined by the distance between anterior-most (top) and posterior-most (bottom) cells. Cells rearrange within the bounding box as it narrows in the ML, driving cell movements in the AP direction until 15% AP strain is reached (Fig 3A, Supplemental Video 4).

**Figure 3.**
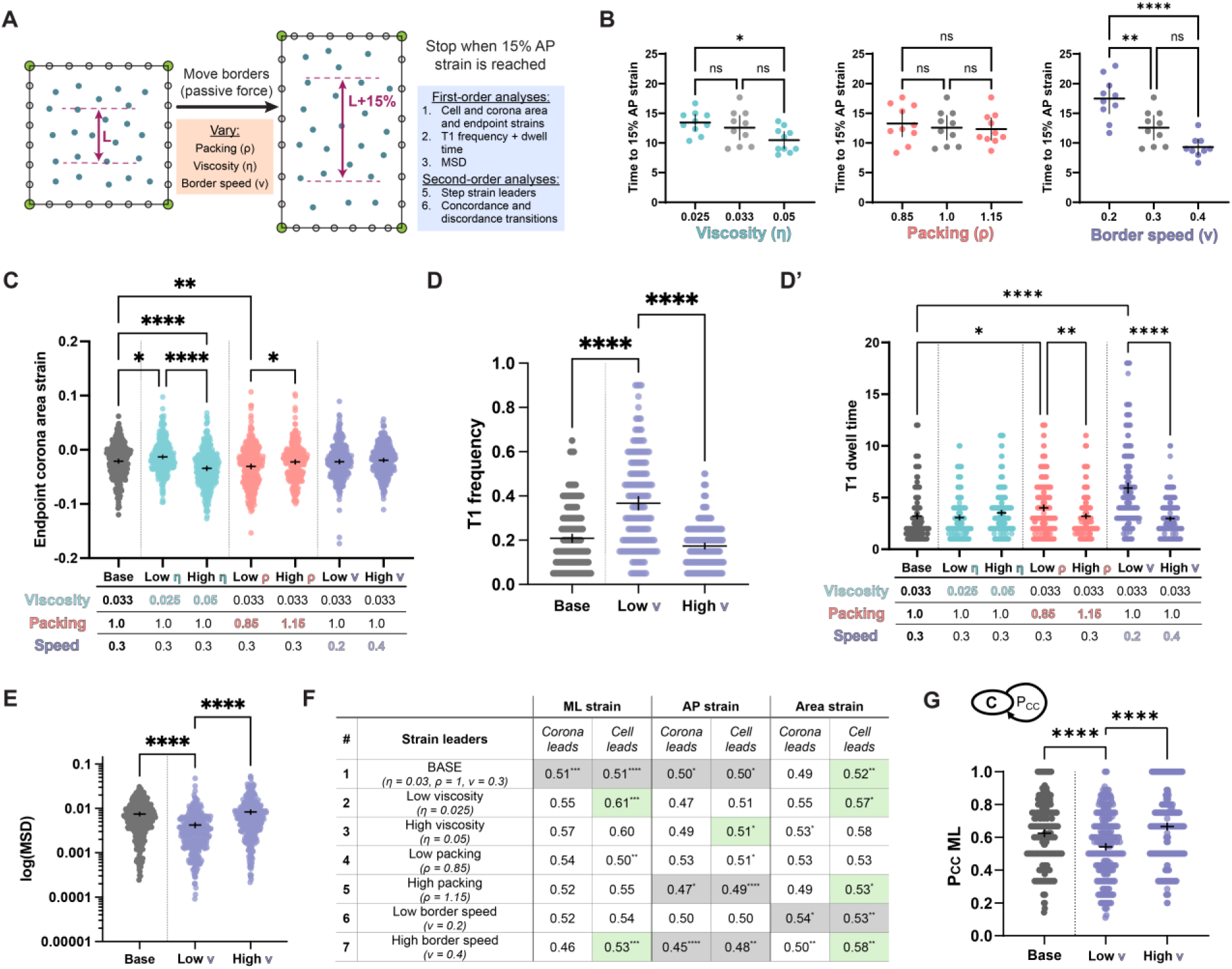
Using the passive model to understand the effects of tissue-wide properties on CE dynamics. [number of cells (n) for each case, 10 simulations per case: (1) 307, (2) 307, (3) 324, (4) 324, (5) 326, (6) 326, (7) 346] (A) Schematic of the application of passive force to simulated cell centroids, which is then tessellated for analysis. (B) Comparison of the number of timesteps it takes for passive simulations to reach 15% AP strain under conditions of changing viscosity, packing, and rate of CE. (n=10 simulations for each case). (C) Comparison of the base case to each of the other cases, and cases with the same viscosity, packing, or CE rate for the Corona Area strain. (D) Comparison of the base case to each of the other cases, and cases with the same viscosity, packing, or CE rate for proportion of cells in T1 transitions on average over all time frames (D’) How long cells spend in the state of a higher order junction on average per simulation. (E) MSD of the base case compared to each of the other cases, and cases with the same viscosity, packing, or CE rate. (F) Table for cell and corona leading behaviors for each of the seven cases. Green shading in the table indicates that the cell is leading. Grey shading represents cases where both cell and corona-led strain lead metrics are greater than no-shift cases, but there is not a statistical difference between the level of cell or corona-led strain. (G) PCC_ML_ compared between base case and each of the other cases, and cases with the same viscosity, packing, or CE rate.

From cases of successful passive CE, we assess the role of the speed of boundary movement (v), the density of cell-packing (ρ), and the viscosity (η). First, we assess how each parameter affects the time it takes for the cells to reach the stop condition of 15% AP strain. We note that a higher viscosity increases a tissue’s resistance to flow and find that higher viscosity increases the time the simulated tissue takes to reach 15% strain. Packing does not have an effect on the time to reach 15% AP strain, while faster border speed decreases this time (Fig. 3B). Of cell and corona, ML, AP, and area strains, only corona area strains differ as parameters are changed (Fig. 3C). Lowering viscosity and increasing packing increases corona area strains. The speed of boundary movement has no effect on the corona area strain, suggesting that other factors, such as cell rearrangement reduce the effect of external forces. Boundary movement speed is the only parameter that has an effect on the proportion of time cells spend in T1 transitions, with slower border movements resulting in more cells participating in T1 transitions (Fig. 3D). Lower border speeds also cause cells to spend more time dwelling in T1 transitions (Fig. 3D’). Lower cell packing also increases the time that cells spend in T1 transitions. This result suggests that cells are locked in transition states and precluded from moving. This appears to be the case, as slower border speeds lower cell MSD (Fig. 3E). Taken together, less cell movement and more cell rearrangements may limit the amount of corona area strain during passive CE.

We applied our second-level analysis to determine relationships between cell and corona strains during passive CE in response to external forces. Lower viscosity and higher border speeds indicate cells leading ML strain, with cell-led strain exceeding ML strain with no leader (Fig 3F, green shading, Supplementary Fig. S4A). However, the base case also results in cells leading ML strain, but this is not significant because a similar level of coronas leading ML strain is observed, indicating no overall strain leader in that direction (Fig 3F, grey shading, Supplementary Fig. S4B). The base case, along with low viscosity, high packing, and high CE speed result in cell-led area strain (Supplementary Fig. S4C). We observe that slower CE speed results in cells spending less time staying in concordance in the ML direction (Fig 3G). This trend is also observed in the AP direction (Supplementary Fig. S4D). This suggests that more rapidly applied external force promotes more coordination between cell and tissue strains. With a description of how cells and tissues change shape as CE is driven by external forces, we investigate cell and tissue shape changes during CE generated by active, force-generating cells.

### Multiple active force-generating cell behaviors can drive CE

Multiple cell behaviors can drive CE in the absence of external forces. To remove the physical constraint of the tissue borders, the border cells are converted to free cells (Fig. 4A). We next introduce an attractive force to maintain tissue cohesivity to counteract the existing repulsive force pushing cells outward. We select a Lennard-Jones potential (Lennard-Jones 1931) function so that attraction force between cells decreases when they are further apart. The attractive force is applied once a cell is further than one cell diameter from the centroid of its neighbor (the repulsion range), and stops at two cell diameters, mimicking cell contacts *in vivo*. The Lennard-Jones potential is integrated to give us force:

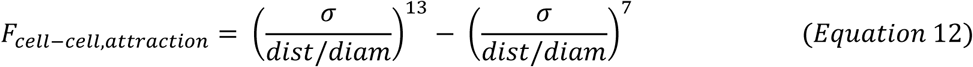

**Figure 4.**
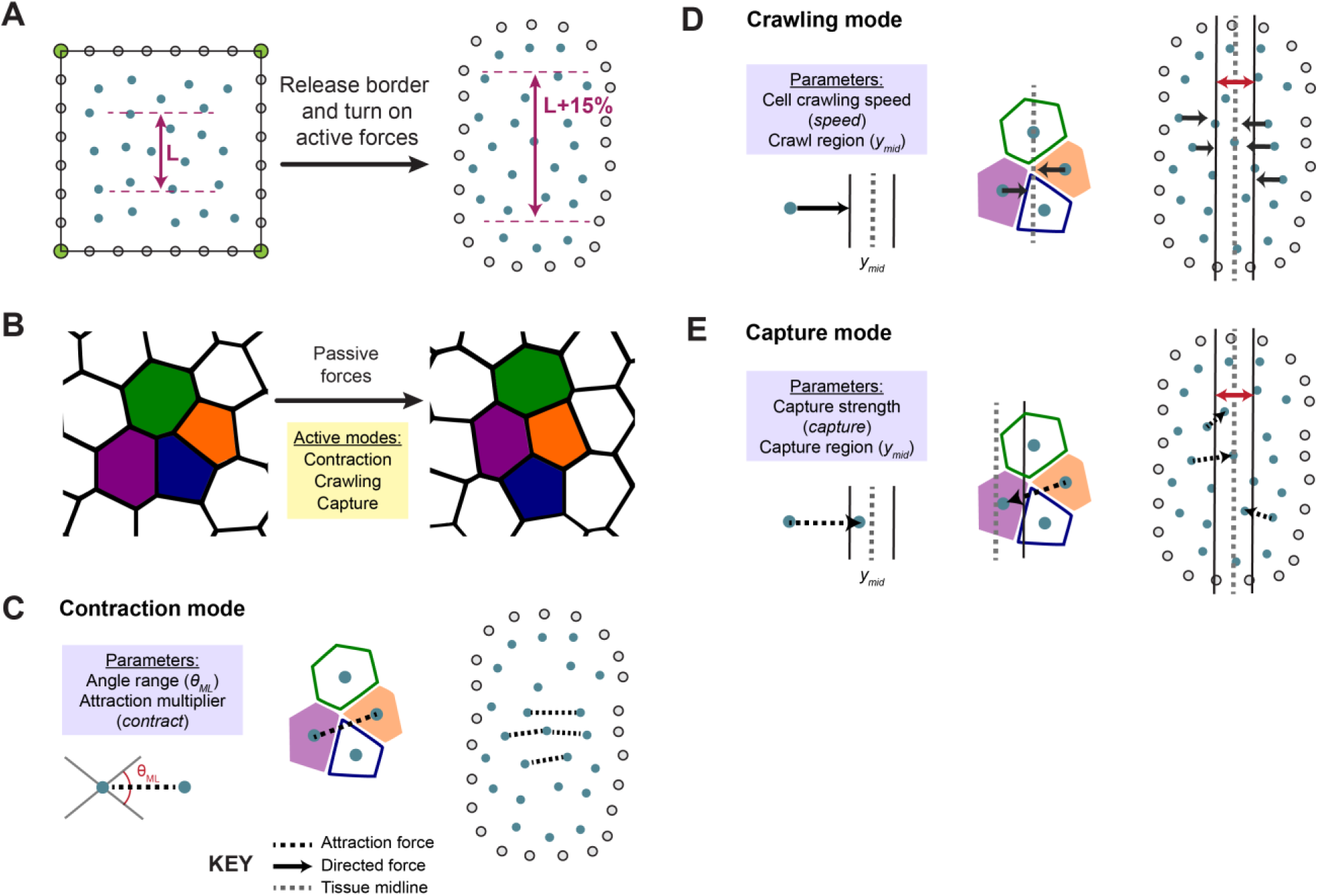
Representing different modes of active-force driven CE in the model. (A) When active forces drive CE in the mode, the border cells are converted to free cells. The simulation still stops once the interior region reaches 15% AP strain. (B) The cell rearrangements characteristic of CE can be achieved independently by the passive mode, or any of three active modes. (C) The contraction mode’s parameters are an angle range in which the attractive force between cells is scaled by a multiplier. (D) The crawling mode’s parameters are the speed by which cells undergo directed motion and a midline region in which they cease the directed motion. (E) The capture mode’s parameters are a range within the midline where cells outside are being acted on by an attractive force that is scaled by a multiplier.

where the first term represents repulsion, and the second term represents attraction. σ is the distance at which the potential is zero, equivalent to rest_ij_ (Equation 3).

To investigate the difference in active and passive cases of CE we investigate three modes of cell behaviors, each capable of driving CE in the same time-frame as the passive case: contraction, crawling, and capture (Fig 4B). The contraction case is characterized by enhanced pairwise attraction within an angle range (θ_contract_) of the ML axis (θ_ML_) via a multiplier (*contract*) applied to F_cell-cell, attraction_ (Fig. 4C, Supplemental Video 5).

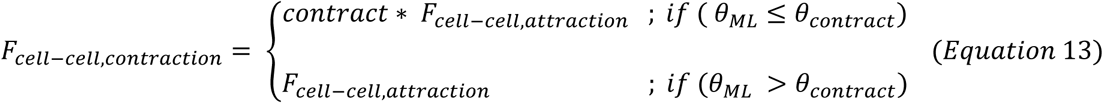

For the crawling mode, cells outside of the midline region of width (y_mid_) exhibit directed movement (*speed*) towards the midline (Fig 4D, Supplemental Video 6).

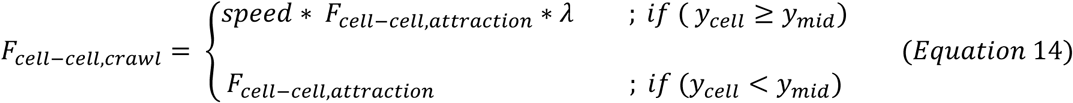

To introduce heterogeneity, a set speed in pixels per timestep is multiplied by a uniformly distributed random number (λ). The final midline capture mode has properties of both contraction and crawling modes. Cells within the midline region exert attraction multiplied by a strength factor (*capture*) on cells that lie outside this midline region (Supplemental Video 7).

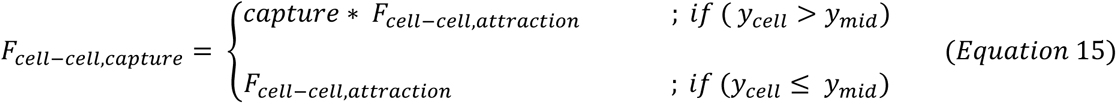

Parameters that regulate each of these three modes can affect the rate of CE: i) for the contraction mode, strength of the contraction and its angular range, ii) for the crawling mode, the range of the attractive signal and the speed, iii) for the capture mode, the range of capture and strength of the attractive force. Parameters were selected that enable a 15% AP strain from the initiation of active forces, like the passive case (Fig 5B, see Supplementary Table 3 for parameters).

**Figure 5.**
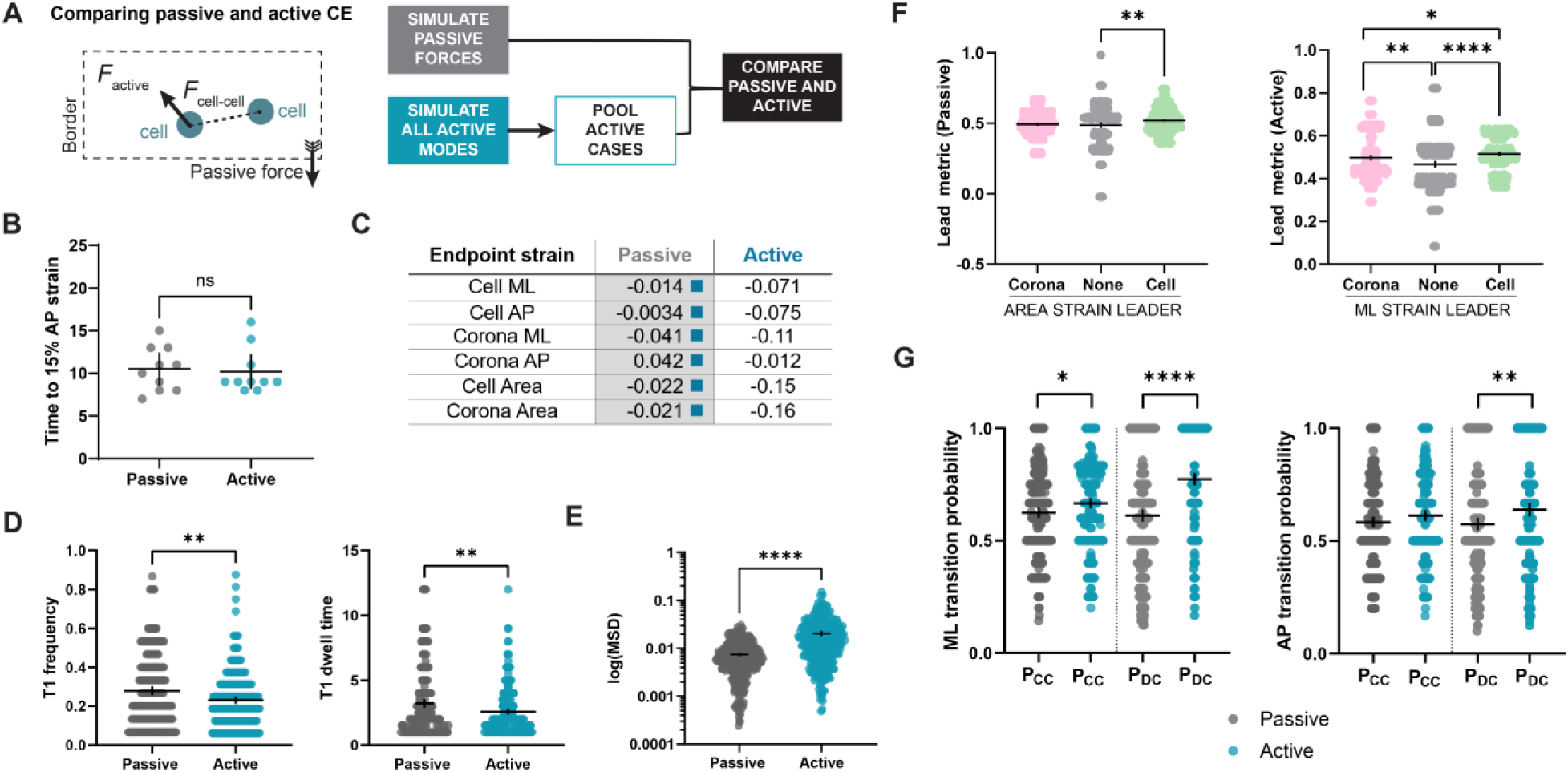
Comparison of the dynamics of the base passive case and a pooled active case. [n= 307 cells for passive across 10 simulations, n=287 for active pooled across 10 simulations]. (A) Schematic showing how active cases are initiated in the model. (B) Comparison of time to 15% strain between passive and active cases. (C) Comparison of cell and corona ML, AP, and area strains between passive and pooled active cases. The grey shading marks the higher mean for significantly significant differences, with the color matching the case that the mean exceeds for a given comparison. (D) T1 proportion and dwell time plots comparing passive and active cases. (E) MSD comparison between passive and active cases. (F) Comparison of cell and corona leading behaviors for passive and active cases for the same groupings as Figure 3. (G) Comparison of transition probabilities for passive and active cases for the same groupings as Figure 3, with plots for all comparisons.

### Comparing passive and active CE

To assess how cell and tissue morphodynamics differ for passive and active CE, we compare the outcomes of our analyses applied to passive and active simulations. Before we proceed to comparing passive CE to each active mode, we compare passive simulations to a set of cases drawn from contraction, crawling, and capture active modes. We refer to this set of active simulations as a “pooled active”, or simply the “active” case (Fig 5A). This approach enables distinction between the effects of passive and active modes regardless of the exact active mechanism driving CE. Comparisons between individual modes will be discussed below.

Unlike the comparisons between the different passive cases where only corona area endpoint strain differs (Fig 3C), there are differences in all endpoint strains between passive and active cases (Fig. 5C, shaded cells indicate the statistically higher mean strain, with the colored box indicated the case it is compared to; Supplementary Figure S5). Simulated cells that undergo CE via active modes exhibit lower endpoint cell and corona ML, AP, and area strains. Active cells undergo 5% fewer T1 transitions than passive cells and spend less time in higher order junctions (Fig. 5D). It has been proposed that T1 transitions indicate the presence of active forces (Choi et al. 2016; Jain, Voigt, and Angheluta 2024), but we do not observe this. Furthermore, active modes exhibit higher MSD than passive modes (Fig. 5E).

Continuing with the second order analysis, we find cells undergoing passive CE lead their area strains. Cells engaged in active modes of CE lead their ML strains (Fig. 5F). Comparing concordance and discordance transition probabilities, in the ML direction, active cells have significantly higher P_CC_ with little difference in the AP direction (Fig. 5G). Furthermore, active cells have higher P_DC_ for both axes (Fig. 5G). Thus, whether the initial state is concordance or discordance, active cells tend to move toward concordance in both the ML and AP axes.

### Distinguishing between different types of active strain

We observe distinct mechanical outcomes for cells undergoing CE due to passive and active forces. We expect that contraction, crawling, and capture modes will yield differences in cell and tissue kinematics (Fig. 6A). We compare each active mode, as well as distinguishing each from the pooled active case. We confirm that the time to 15% AP strain is consistent across active modes (Fig. 6B). We find that cells undergoing CE via the contraction mode have higher cell ML, cell AP strain, corona ML, cell area, and corona area endpoint strains than the other cases, including the active pooled case (Fig. 6C, full data in Supplementary Figure S6). Cell and corona ML and area strains are reduced for the crawling case as compared to active pooled cells. The capture mode also has a lower corona area strain than the active pooled cells. There is no significant difference between corona AP strains across all modes, suggesting endpoint strain differences between active cases are manifested predominantly in the ML direction. Cell area strains are significantly different from the pooled active case only in the crawling mode.

**Figure 6.**
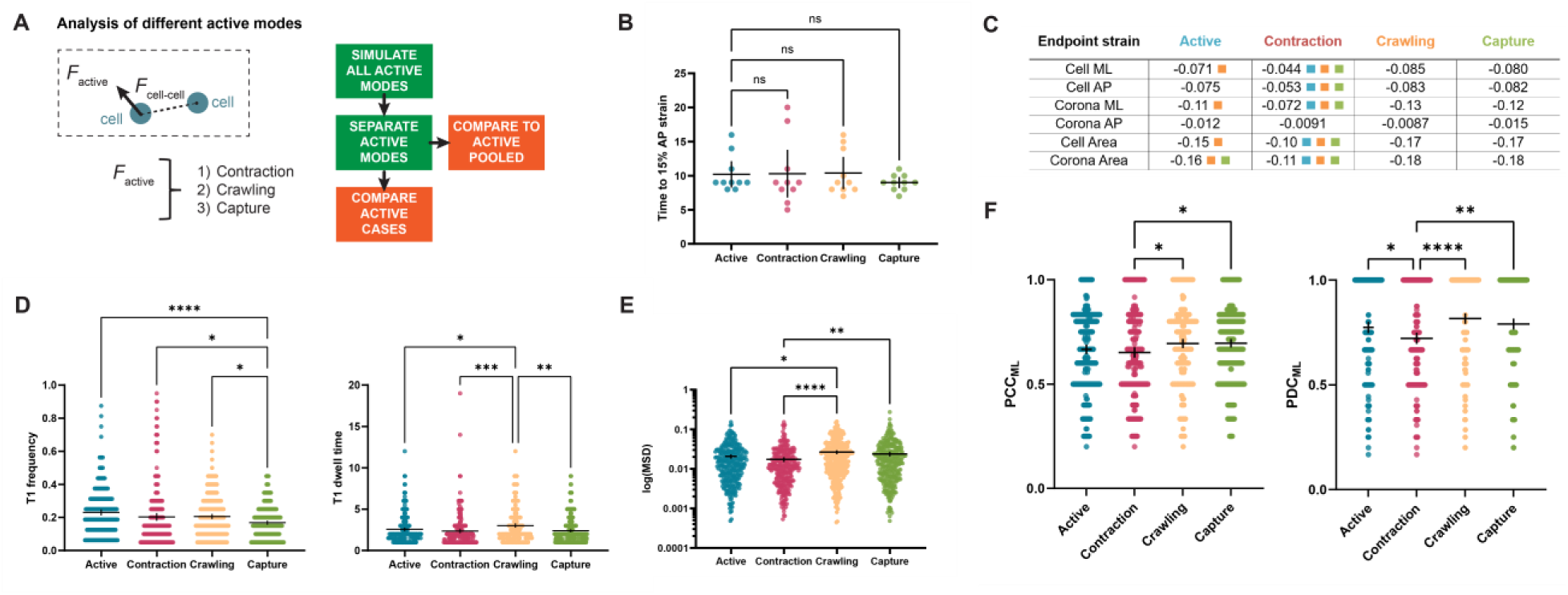
Comparing different modes of active-force-driven CE. (10 simulations per case, n=287 cells for active pooled (same data as Figure 5), n=292 (Contraction), n= 305 cells (Crawling), n=305 cells (Capture)] (A) Diagram of analysis rationale differentiating Figure 6 from Figure 5. (B) Comparison of time to 15% AP strain for the active pooled case and the three active modes: contraction, crawling, and capture. (C) Comparison of endpoint strains for the active pooled case and the three active modes. The values shown are the means, and the colored squares indicate which mean a given mean is significantly higher than. For example, the orange box next to the first mean in the table indicates that the cell ML strain in the active pooled case is significantly greater than only the crawling case. (D) T1 frequency and dwell time comparison for the active pooled case and the three active modes. (E) log(MSD) comparison for the active pooled case and the three active modes (F) Transition probability comparison in the ML direction for PCC and PDC for the active pooled case and the three active modes.

The lowest number of T1 transitions are in the capture mode, at a level significantly lower than the pooled case and all other active modes (Fig. 6D). T1 dwell time is highest in the crawl mode as compared to all other cases (Fig. 6D). The MSD of the contraction mode is the lowest and significantly different from the capture and crawling modes, but not from the pooled active cells (Fig. 6E). The crawling mode’s MSD is higher than the pooled active cells. Leading behavior is not significantly different between modes (data not shown). Concordance probabilities only differ in the ML direction between active modes, with crawling and capture modes showing the greater trend toward concordant states (Fig. 6F). There are no significant differences between transition probabilities in the AP direction (data not shown). The many differences in cell and tissue behaviors in the contraction, crawling, and capture modes can allow us to discern whether these modes are present *in vivo* during CE.

### Assessing passive and active modes of CE across the *Xenopus* neural plate

Having characterized simulated modes of CE, we compare these modes to CE in the *Xenopus* neural plate. The posterior neural plate undergoes 15% AP strain between stages 12 and 15, during which 15% AP strain is reached over one hour (Fig. 7 A-A’). To assess spatial differences across the neural plate, we binned our data into three regions: midline, lateral 1, and lateral 2, with lateral 2 cells being the most distant from the midline (Fig. 7B). We have also pooled all regions to represent the full width of the neural plate. We observe differences in both T1 frequency and duration across the neural plate. The midline region experiences the highest T1 frequency and duration, which may suggest that the midline region is passively responding to external forces (Fig. 7C). The most lateral region has the fewest T1 transitions and lowest dwell time, suggesting the involvement of active modes. Furthermore, the midline exhibits the lowest MSD and the most lateral region the highest, also suggesting the involvement of active modes (Fig. 7D).

**Figure 7.**
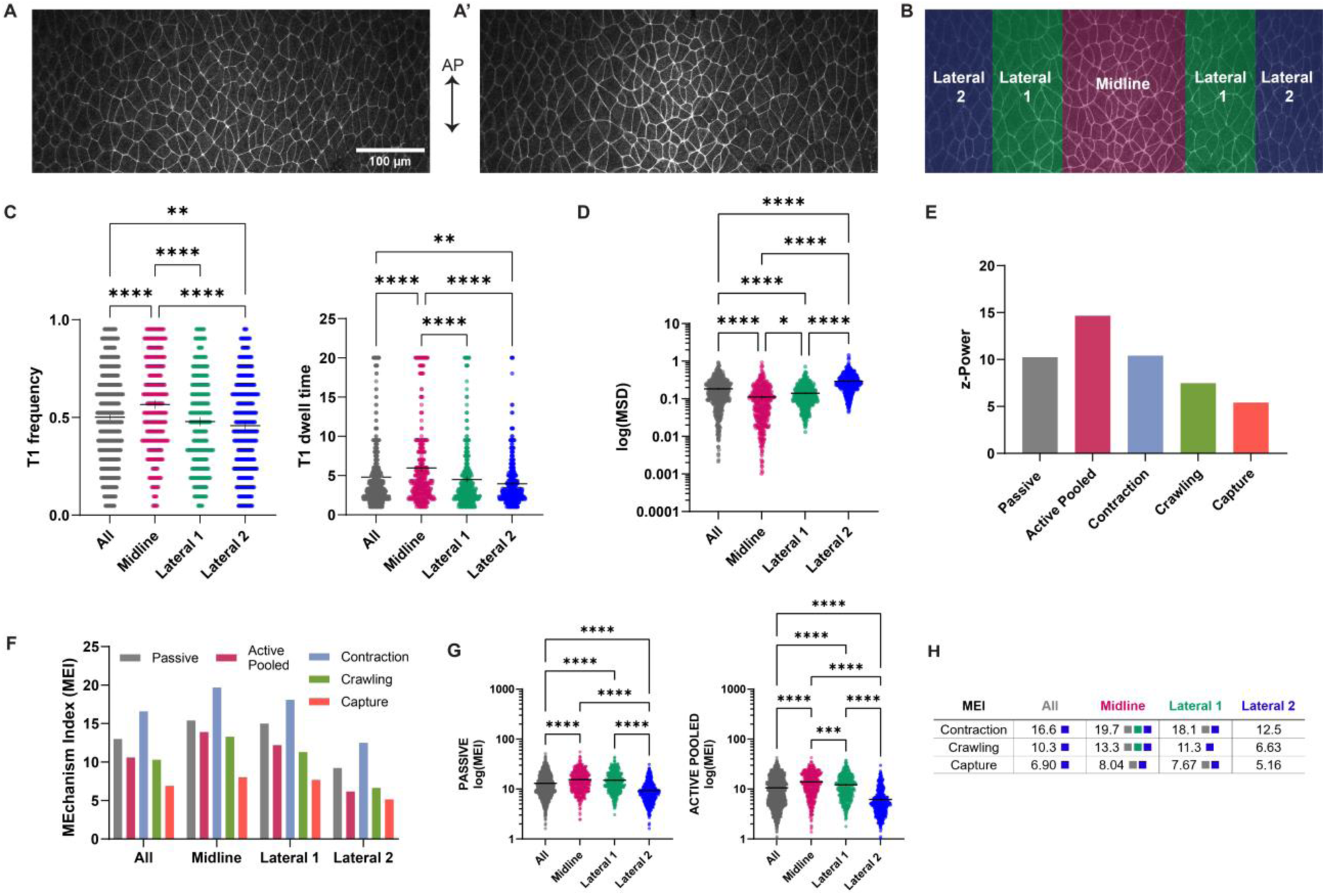
Applying mechanism inference to live-cell CE across the Xenopus neural plate. [frog n =1354 cells across 3 explants and 2 whole embryos, n = 442 cells (midline), n = 432 cells (lateral 1), n = 480 cells (lateral 2)] (A) *Xenopus* neural plate at stage 12 as neural CE begins. (A’) *Xenopus* neural plate at stage 15 as neural CE continues and 15% AP strain is reached. (B) Division of the neural plate into three regions to assess spatial variations-the colors scheme is used throughout the figure. (C) Comparison of T1 frequency and dwell time between the entire region shown and the midline, lateral 1, and lateral 2 regions. (D) Comparison of MSD between the entire region shown and the midline, lateral 1, and lateral 2 regions. (E) z-Power, or sum of z scores distinguishing each case for mechanism inference. (F) MEI for the entire region shown and the midline, lateral 1, and lateral 2 regions. (G) Passive and active pooled MEI compared between the entire region shown and the midline, lateral 1, and lateral 2 regions. (H) Contraction, crawling, and capture MEI compared between the entire region shown and the midline, lateral 1, and lateral 2 regions.

### Developing a quantitative framework for mechanism inference

We seek to directly quantify the similarity between CE in live cells and simulated cells in the modes compared. Given that we have several metrics for each case simulated, each with different means and degrees of variation, normalizing the data enables more consistent comparisons. The magnitude of z-score differences has been used to distinguish expression profiles in genomic datasets (Thomas et al. 2001; Webb-Robertson et al. 2011). To this end, the z-score of each significantly different metric from the simulations for each case relative to one or more reference cases is calculated (see supplemental text). This provides intuition of the difference in mean and standard deviation of our first and second-order analyses. ‘Z-power’ is the sum of z-scores differentiating a given case from its reference case(s). A z-power of 15.47 indicates that a case is 15.47 standard deviations away from its reference case in total. Using the results of our first and second order analyses, the passive case’s reference case is the active pooled case. Similarly, the contraction case’s reference cases are the active pooled case, the crawling case, and the contraction case (Fig. 7E). Z-scores that are significantly different from the reference case (*i* through *n*) are used to calculate the MEchanism Index (MEI) (see supplement):

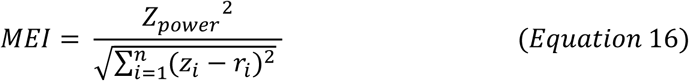

We apply this analysis to CE across the *Xenopus neural plate*, as well as the midline, lateral 1, and lateral 2 region. We see that the midline region has the highest passive MEI, supporting our observation of higher T1 frequency and dwell time, as well as lower MSD (Fig. 7F-G). In addition, we see that the midline has the highest MEI for the active pooled and individual active cases. To explore this further, we compared each active mode between the full neural plate and each region (Fig. 7H, full data in Supplementary Figure S7). The lateral 2 region has the lowest MEI compared to all modes, followed by the lateral 1 region. The lateral 1 region exhibits more contraction and capture behaviors than the neural plate as a whole. This suggests that certain active modes of CE may be less distinguishable as distance from the midline increases or that the modes that we model are not fully representative of the behaviors in the lateral 2 region.

### Conserved inferred mechanisms of CE across developmental model species

As CE is a highly conserved process, we sought to assess how the inferred forces from our analyses would differ across species. CE occurs in the fruit fly *Drosophila melanogaster* during germ band elongation (GBE) and in the mouse *Mus musculus* during neural plate elongation (Supplemental Video 8). The fly germ band undergoes 15% strain from the start of imaging at stage 7 (Fig. 8A) and the end of imaging at stage 8 (Fig. 8A’). By contrast, the mouse neural epithelium has a much lower strain rate due to its slow rate of development (Fig. 8 B-B’, Supplemental Video 9). Due to the lower overall strain, mouse neural data may limit the inferential power of our second-order analyses. We contrast the CE in fly and mouse to CE in the midline of the frog neural epithelium, as these regions similarly undergo the most strain. We use the same number of time lapse frames for all model organisms.

**Figure 8.**
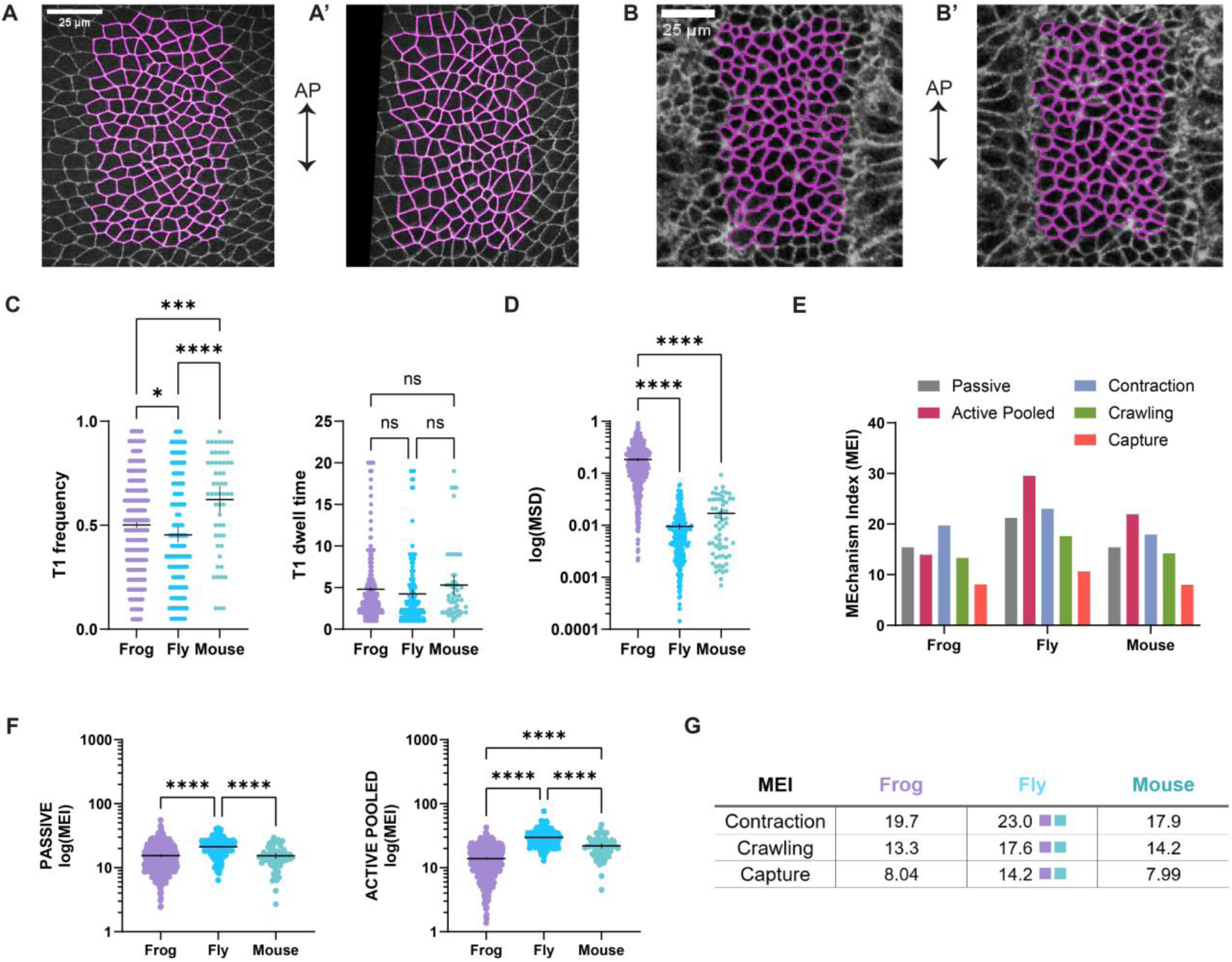
Comparing conserved inferred mechanisms of CE across model species. [n = 227 cells across 5 samples (fly), n = 62 cells across 2 samples (fly)] (A) Segmentation from representative fly embryo at the beginning of GBE (stage 7), labeled with ubi-E-cadherin:GFP. (A’) Segmentation from representative fly embryo after 15% strain is achieved along the anterior-posterior axis. (B) Segmentation from representative mouse embryo at the beginning of neural CE (E8.0), labeled with mEGFP. (B’) Segmentation from representative mouse embryo later in CE (E8.25). (C) Comparison of T1 frequency between frog, fly, and mouse embryos undergoing CE. (D) Comparison of MSD between frog, fly, and mouse embryos undergoing CE. (E) MEI for each class of mechanism inference for frog, fly, and mouse embryos undergoing CE. (F) Passive and active pooled MEI compared between frog, fly, and mouse embryos undergoing

When comparing T1 transition frequency and dwell time across the three species, we find fly cells undergoing CE exhibit the lowest T1 frequency, while the mouse cells have the highest (Fig. 8C). There are no significant difference in T1 dwell times across species. Frog cells have the highest MSD, followed by mouse, and fly (Fig. 8D). From the MEI values, it appears that passive behaviors are most common in fly cells (Fig. 8E). For active pooled behaviors, fly CE matches the simulation reference datasets most closely. Frog cell MEI is higher for active pooled behaviors than in mouse, possibly due to the lower rate of CE in mice. Fly CE cells experience higher MEI for all active modes compared (Fig. 8G, full data in Supplementary Figure S8). The higher T1 frequency and lower MSD for mouse cells suggests that passive CE may be a better explanation for the MEI values than active behaviors. The MEI values for all active modes are not significantly different between the neural vertebrate cases of frog and mouse, despite differences in the rate of AP tissue elongation. Comparison of MEI values in tandem with descriptive analyses across model species allows us assess conserved mechanisms of CE.

## DISCUSSION

Cells form tissues and organs during morphogenesis via mechanical integration of cellular force-generating machinery (Davidson 2024). Convergent extension (CE) describes the tissue deformations from varying sources of active force generation, but CE can also occur as a passive movement in response to forces generated outside the tissue of interest. Thus, when tissues are observed to undergo CE, we cannot assume it is due to any specific active force or even that motions are self-organized. While CE has been observed in a broad range of developing tissues (Keller et al. 2000; Warga and Kimmel 1990), few instances have been the subject of detailed study (Weng, Huebner, and Wallingford 2022; Tada and Heisenberg 2012). The work described here is intended as a framework for using simulations to distinguish passive from actively-driven CE, in addition to providing a way to quantify differences between specific mechanisms in live-cell data. We refer to this process as “mechanism inference”.

We developed a series of computational active particle models of epithelial CE. We have two main classes of models: 1) a passive model, representing CE in response to external forces, where tissue borders move to drive CE, and 2) three active models where a localized contraction mode, a crawling mode, and a capture mode drive CE. In each of these, we represent the pattern of force generated by the actomyosin cytoskeleton of active cells via a distinct force profile based on observations in live tissues (Weng, Huebner, and Wallingford 2022; Gorfinkiel and Blanchard 2011; Sutherland, Keller, and Lesko 2020). It has not been described how each of these mechanisms differentially affects tissue morphometrics *in vivo*. We leverage a suite of analyses to characterize our simulations and compare them with live cell data to infer mechanisms contributing to live cell CE.

We divide our characterization into first-order and second-order analyses. The first-order analysis consists of endpoint strain data in both tissue axes and area for a cell and its local neighborhood (corona), as well as quantifying the frequency and duration of T1 transitions, and the MSD of cells. Our second order analyses are based on step strains, or strains between subsequent time frames of simulations or live cell time lapses. In our strain-leader analysis, we quantify whether a cell or its corona is initiating strain, and in our concordance analysis, we assess whether cell and corona strains are coordinated as they change over the course of CE.

We first characterize cell and tissue dynamics in the purely passive model and compare the effects of the speed of tissue boundary movement, cell packing density, and tissue viscosity and report the differences in our first and second order analyses. Our findings suggest that lowering tissue viscosity reduces corona strain, and the opposite occurs if there is more resistance to movement. Both higher border speed and lower viscosity cause cells to lead strain in the ML direction, suggesting that higher viscosity and lower border speeds dampen ML deformation. We compare simulations of passive CE to a pooled active case consisting of samples from all three active modes to discern which of our analyses may distinguish passive and active CE. We find that passive models undergo more frequent T1 transitions and spend more time in higher order junctions. This observation contrasts with conventional views of CE where T1 transitions are attributed to sites of active force generation. Passive cells lead their corona area strains and have greater endpoint strains overall, while active cells lead their corona ML strains. Our results suggest that cells undergoing CE in response to external forces experience more strain over time and respond by remaining in T1 transition states, as they do not lead their coronas in ML direction strain. Moreover, passive cells are less likely to remain in concordance with their coronas. This further implies that T1 transition processes allow cells to “resist” strain imposed from their immediate neighbors. Cells undergoing CE via the capture mode exhibit the lowest T1 transition frequency, while cells in the crawl mode spend the most time in T1 transition states. Contracting cells have the lowest displacement, suggesting that bipolar anisotropic tensions offset the effects of local displacements. Cell and corona strains are more concordant in crawling and capture modes, which may reflect the ability of these modes to counteract higher displacements.

To infer mechanisms of CE *in vivo*, we consider early movements of the posterior neural plate in *Xenopus*. We observe more T1 transitions, higher dwell times, and lower MSD in the midline compared to lateral regions, suggesting more passive behaviors. To establish a framework for inference, we create reference datasets to derive a ‘z-power’ to normalize differences in passive, active pooled, contraction, crawling, and capture modes. Within this framework, we introduce a MEchanism Index (MEI) to compare CE *in vivo* to reference cases of each simulated mode of CE. Comparing frog CE analyses at various distances to the midline of the embryo, we find midline cells are more similar to passive modes and also more similar to active modes than more lateral regions. We find higher MEI for the lateral 1 region for contraction and capture modes compared to the neural plate a whole. Lower MEI at greater distances from the midline may suggest CE is driven by mixed modes with high cell-cell variability in more lateral regions. Finally, we explore conserved mechanisms of CE by comparing inferred mechanisms from the frog midline to both fly and mouse CE. This comparison suggests that fly cells are undergoing CE more by discernable active modes than frogs or mice. Furthermore, similar MEI values for frog and mouse for all active modes confirm conserved mechanisms in vertebrate neural CE in particular. These differences have not been reported previously, and are an exciting avenue for continuing research.

The results from this study can inform experimental design by focusing attention on spatial patterns of cell and tissue shape changes produced by each active mode. *Xenopus* cells can be injected with factors that specifically perturb contraction, crawling, and capture or extend these behaviors to more lateral regions. Due to the dependence of contraction, crawling, and capture approaches operating using similar contractile machinery, it may be promising to apply optogenetic or photoactivation approaches (Cavanaugh et al. 2020; Oakes et al. 2017). The activation or inhibition of contractility at tricellular junctions where protrusions form may also reveal how force is transmitted. Tissue-wide properties of viscosity and packing can be modified experimentally via modulating calcium flux and microtubule inhibition (de Leeuw et al. 2024; Bera et al. 2022). Assessing what morphological changes these manipulations drive in epithelial tissues may open further avenues of inquiry.

Several limitations of our study emerge from implicit modeling assumptions. A limitation of our study is that basal contributions to CE are not considered. Basal-first cell rearrangement is a current topic of study (Sun et al. 2017). Moreover, the *Xenopus* neural plate consists of both an epithelial layer and a mesenchymal layer. Cell rearrangements can propagate in multiple planes and integrate into observed apical cell shapes, indicating a passive response. Several active modes that we test in this paper, crawling and capture, are also observed in mesoderm cells (Pfister et al. 2016; Davidson et al. 2006; Elul and Keller 2000). The effect of basal and mesoderm cell perturbations on the inferred modes of CE may yield insights into the integration of force production and mechanosensation across the embryo.

In summary, we developed techniques to simulate distinct modes of CE and a framework to infer mechanisms from live-cell data via a robust statistical engine. The findings from this study can guide experimentation and further our understanding of the forces shaping tissues and organs during morphogenesis.

## METHODS

### Live cell experiments

#### Xenopus laevis (frog)

*Xenopus laevis* embryos were obtained by standard methods (Kay 1991), fertilized *in vitro*, and cultured in 1/3X Modified Barth’s Solution (MBS) (Sive 2000). For live visualization of the apical surface, mRNA-encoding fluorescent protein constructs were injected at the one-, two-, or four-cell stage to visualize the plasma membrane (CAAX-GFP, CAAX-mCherry). For explants, embryos are cultured to stage 12.5 (Nieuwkoop and Faber 1956), at which time dorsal axial and paraxial tissues (dorsal isolate) are microsurgically removed from the embryo using hair loops and knives in DFA solution (Danichik’s For Amy; (Sater, Steinhardt, and Keller 1993)). Before microsurgery, small (∼1×5mm) pieces of glasses are cut from a #1.5 glass slide using a diamond pen. After isolation, tissue explants are transferred to a clean dish filled with DFA and gently and minimally compressed under a precut small piece of glass using vacuum grease at the ends for half an hour prior to experimentation to allow healing without tissue folding/bending. For microinjection of mRNAs, embryos are placed in 1X MBS containing Ficoll. Imaging was performed on a spinning disk confocal (Yokagawa CSU-X1) using a 25x Water Objective to produce tiled z-stacks (2μm, Z-step size) images of the tissue. Confocal stacks were collected at 3 minute time intervals. We imaged apical surface of dorsal isolate explants held under a glass coverslip or of whole embryos mounted in 1% ultra-low melting agarose (type IX-A; Sigma) gels in custom imaging chambers. All frog animal use protocols were in compliance with PHS and USDA guidelines for laboratory animal welfare and reviewed and approved by the University of Pittsburgh Institutional Animal Care and Use Committee.

#### Drosophila melanogaster (fly)

*Drosophila* embryos expressing ubi-E-cadherin:GFP (Oda and Tsukita 2001) were collected for 1 hour, dechorionated in 50% bleach and transferred to a drop of halocarbon oil 27 on a piece of cover glass (Sigma). Embryos were mounted on an oxygen-permeable membrane (YSI). We used a 63X oil immersion lens (Zeiss, NA 1.4) to image embryos on a CSU-22 spinning disk confocal microscope (Yokogawa). We collected 16-bit z-stacks at 0.5 μm steps every 15 seconds (15 slices/stack). Maximum intensity projections were used for analysis.

#### Mus musculus (mouse)

Natural matings of mTmG-Sox2Cre mice were set up and checked daily for the presence of a vaginal plug indicating mating. The morning a plug was identified was designated as E0.5 of pregnancy, and embryos were dissected at E8.0 – E8.25 in Whole Embryo Culture Medium (WECM) (Yen et al. 2009). Embryos were then cultured in a mixture of 50% WECM and 50% rat serum. For imaging, embryos were mounted distal tip down in a glass-bottomed chamber, covered with culture medium, overlaid with mineral oil to prevent evaporation, and imaged using a Zeiss 780 or Leica SP8 confocal microscope in an environmental chamber at 37°C and 5% CO_2_ gas. For CE and cell behavior analysis, embryos were imaged for 6-10 hours at an interval of 6min at 40x with Z-stacks of 10-15 planes captured at 2μm intervals. All mouse animal use protocols were in compliance with PHS and USDA guidelines for laboratory animal welfare and reviewed and approved by the University of Virginia Institutional Animal Care and Use Committee.

### Image analysis

*Xenopus* fluorescent membrane outlines were segmented using SeedWater Segmenter, a semiautomated watershed algorithm based segmentation software (Mashburn et al. 2012). Cellpose (Stringer et al. 2021) was used within Track Mate (Ershov et al. 2022) in FIJI (Schindelin et al. 2012) to segment fly and mouse cells. The binary cell outline and cell identity maps outputted by the segmentation programs were processed with custom written FIJI macros to extract individual ROIs for each cell. Cells from the centers of segmented regions are used for analysis to allow for complete coronas. Further analysis quantifying cell morphology parameters, cell neighbor exchanges, and strain was performed in MATLAB (MathWorks).

### Simulation and analysis pipeline

The simulation, as described in our previous paper, is run via an integration between FIJI and MATLAB (Sage et al. 2012; Anjum et al.). The simulation parameters and file paths are set in MATLAB and read-in by FIJI when the centroids are moved based on our force balance. These are saved as regions of interest (ROIs) which are read-in and used to perform a Voronoi tessellation. For automated segmentation of simulation results, the tessellation is passed through the “Analyze Particles” routine in FIJI with ROIs of each cell being filled in with their unique ID from the simulation. This enables us to track the dynamics of each cell over time. The simulation starts with equilibration, followed by moving the boundaries so that the tissue is in a dynamic state once we perform our measurements (see supplementary text and Supplementary Figure S2). Simulations are sampled for analysis by downsampling frames by a factor of twenty to match live cell time steps and reduce computing time for analyses. Simulation results are exported to MATLAB to conduct analysis with custom code.

### Strain calculation from ellipses for cell and corona

The deformation gradient matrix F is derived from the equation of an ellipse fitted to the ROI of a cell or its corona and is used to calculate Lagrangian finite strains.

We define the initial configuration of the ellipse (fitted to either the cell or corona) at t=0 as:

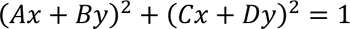

The same ellipse after deformation, at t=1 is defined as:

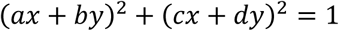

Using the deformation gradient tensor **F**, we map the initial configuration of the ellipse to its current configuration and solve for **F**:

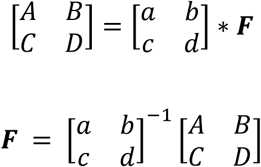

We then calculate the Lagrangian finite strain tensor for both the cell and corona levels.

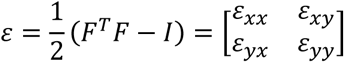

The *xx* strains correspond to ML strains and the *yy* strains correspond to AP strains.

### Markov analysis

For our states of concordance and discordance, we wanted to further quantify the transitions of cells between these states over our timelapses. One way to do so is a two-state Markov model. In this model, there are the two states of concordance and discordance (Fig. 2B-C). We calculate the state transition probabilities i.e. use the time series data to determine the probability of cells to undergo the following transitions:

1. P_DD_: The probability of a cell in a discordant state with its local tissue to stay in this discordant state in the next time step
2. P_DC_: The probability of a cell in a discordant state with its local tissue to move to a concordant state in the next time step
3. P_CD_: The probability of a cell in a concordant state with its local tissue to move to a discordant state in the next time step
4. P_CC_: The probability of a cell in a concordant state with its local tissue to stay in this concordant state in the next time step

The state transition probability is calculated for each transition as follows:

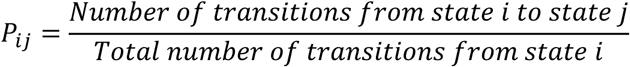

These probabilities are assembled into the state transition probability matrix:

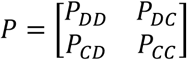

The transition probability matrix **P** was calculated using step strain data for every cell analyzed for every timestep in the simulation for which its strain was calculated.

### T1 transition detection and analysis

T1 transitions are identified by passing a circle with a radius of seven pixels over all coordinates *x,y* of each segmented image. If the number of unique intensities in the circle, excluding the background and cell-boundaries, exceeds three, the identities of these cells and the time are stored. This data is used to populate a matrix of zeroes and ones of dimension cells by time to calculate T1 frequency and dwell time, with a ‘1’ indicating that a given cell was participating in a higher order junction at a given time. T1 frequency for each time frame is calculated by taking the average of ‘1’s for cells participating in T1 transitions in each time step. T1 dwell time is calculated by counting the number of ‘1’s in a row for each cell and averaging over the total number of instances that one or more ‘1’s appeared in the T1 matrix for each cell.

### Mean square displacement (MSD) calculations

MSD is calculated as follows for *i* through *n* particles relative to initial positions x(0) and y(0):

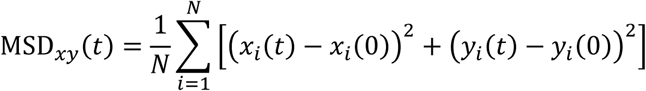

This is normalized by dividing by the cell area (square pixels), resulting in a unitless quantity.

### Statistical methods

Statistical analysis was performed using GraphPad Prism (GraphPad) and MATLAB (MathWorks). Sample sizes are included in figure captions. Simulation number was selected by comparing significance of serial downsampling and matching to live cell data sample size for optimal statistical power (Fig. S3) The significance levels for all figures are indicated with asterisks: * = p<0.05, ** = p<0.01, ***=p<0.001, ****=p<0.0001. All tests are performed at the 95% significance level, with plots showing mean and 95% confidence intervals. We compared area and sidedness distributions with a two-sample Kolmogorov Smirnov test, where higher p-values indicate more similarity between distributions. We summed p-values to account for total variability in setting our base case. To compare two groups, we used two-sample t-tests, and to compare three or more groups for different active modes and live cells, we used one-way ANOVA comparisons. All data was tested for normality and equal variance and confirmed to have sample sizes sufficient to be robust to variation. Post-hoc testing for ANOVA comparisons was done using Šidác’s multiple comparisons test using multiplicity-adjusted p-values.

## Acknowledgements

We would like to thank members from the Davidson lab for their support and helpful discussions. We also thank Adam Kruchten for advice on using the Markov model. This work was supported by grants from the National Institutes of Health, R01 HD044750, R37 HD044750, and R21 HD106629 to LAD. Additionally, SA and DV were supported by the Biomechanics in Regenerative Medicine (BiRM) Training Grant from the NIBIB (T32 EB003392). RFG is funded by the Natural Sciences and Engineering Research Council of Canada (418438-13) and the Canadian Institutes of Health Research (186188). RFG is the Canada Research Chair in Quantitative Cell Biology and Morphogenesis. For image acquisition of mouse embryo neural development we acknowledge the Keck Center for Cellular Imaging at the University of Virginia for the usage of the Zeiss 780 microscopy system (PI: Ammasi Periasamy; NIH-ODO16446) and the Leica SP8X microscopy system (PI: Ammasi Periasamy; NIH-RR025616), and grant support to AS from Eunice Kennedy Shriver National Institute of Child Health and Human Development (R01HD087093).

## Author contributions

SA: Conceptualization, Software, Writing-Original Draft; DV: Investigation, Validation; RFG: Writing-Review & Editing, Project Administration, Supervision, Resources; AS: Writing-Review & Editing, Project Administration, Supervision, Resources; LD: Conceptualization, Writing-Review & Editing, Project Administration, Methodology.

## Materials availability

Not applicable.

### Data and code availability

Data supporting the findings presented can be found within the body of the paper and the supplementary text and videos. Additional information and code for the computational model have been deposited in a Dataverse repository. Anjum, Sommer, 2025, “Code for “Inferring active and passive mechanical drivers of epithelial convergent extension””, https://doi.org/10.7910/DVN/3FHYMJ, Harvard Dataverse, DRAFT VERSION

