## Supplementary text, tables, and figures for "Inferring active and passive mechanical drivers of epithelial convergent extension"

### SUPPLEMENTARY INFORMATION

***Initializing model and pre-straining simulated epithelium***

***Boundary conditions for active CE***

***Z-power and MEI for mechanism inference***

---

**Figure S1.** *Area and sidedness*, supporting Figure 1.

**Figure S2.** *Simulation initialization and pre-strain application*, supporting Figure 1.

**Figure S3.** *Data pooling and sample size selection for live cell data and simulations*, Methods.

**Figure S4.** *Passive case sensitivity,* supporting Figure 3.

**Figure S5.** *Passive case endpoint strains*, supporting Figure 5C.

**Figure S6.** *Active case endpoint strains*, supporting Figure 6C.

**Figure S7.** *MEI for active modes by cell for frog data for all regions*, supporting Figure 7F,H

**Figure S8.** *MEI for active modes by cell for frog, fly, and mouse data*, supporting Figure 8G

---

**Table S1.** *Comparing matching of base case candidate parameters to live cell data*

**Table S2.** *Base case parameters for passive case*

**Table S3.** *Base case parameters for active cases*

**Table S4.** *Mean and standard deviations for all metrics for all simulated cases*

**Table S5.** *Z-power for all cases*

---

**Video S1.** Xenopus *brightfield microscopy timelapse of neural plate convergent extension*

**Video S2.** Xenopus *confocal microscopy timelapse of neural plate convergent extension*

**Video S3.** *Simulated centroid timelapse for passive mode*

**Video S4.** *Simulated passive case*

**Video S5.** *Simulated contraction mode*

**Video S6.** *Simulated crawling mode*

**Video S7.** *Simulated capture mode*

**Video S8.** *Fly germband elongation convergent extension*

**Video S9.** *Mouse neural convergent extension*

***Initializing model and pre-straining simulated epithelium***

In our simulations of different force profiles driving passive and active CE, it is important for all simulations to start from the same "base state" so that our results are comparable. The simulation is initialized as a square field in which free cell positions and potentials are initially sampled from a random distribution. The simulation is advanced in time, with free and border cells redistributing in space to reach an approximate state of equilibrium after 100 timesteps (Fig. S2). When the neural plate is imaged in our timelapses, we are capturing a portion of neural CE—the tissue has already undergone shape change, and we typically capture 15% strain from the beginning to end of imaging. In total, 30% strain is achieved. To replicate this in our simulations, we decided to apply a pre-strain to our simulated epithelium. After equilibration, the borders are moved such that the field goes from 0-15% strain in the AP direction. To limit edge effects from the corner and boundary cells and to have complete coronas for analyzed cells, we limit our analysis to the middle third of the field.

***Boundary conditions for active CE***

Many considerations need to be accounted for in order to represent moving boundaries in simulations. For the passive model, to ensure that force per unit length is consistent while the borders are moved, a new resting potential is calculated at each time step for the border cells. For left and right borders, the new rest potential is based on the change in length of the field, while the top and bottom border rest length is set based on the change in width of the field. The scale remains consistent with the original rest lengths from the equilibration phase.

$$rest_{left,right} = {}^{length_{new}}/_{length_{old}}$$

$$rest_{top,bottom} = {}^{width_{new}}/_{width_{old}}$$

Our active model requires adjustments so that cells do not spread too thin or contract too much. One solution is in a vertex model developed with an active boundary with its own elastic mechanics (Barton, Henkes, Weijer, and Sknepnek 2017). However, in our model, when borders are removed, depending on the arrangement of border cells, surface tension dominates and drives tissue rounding, disrupting anisotropic force application. To prevent this, we convert border cells to “free” cells. In the limit of our pairwise attractive force, this allows cells to adhere. The rest lengths remain unchanged from their state at the end of the pre-strain phase so that no additional forces are introduced.

***Z-power and MEI for mechanism inference***

We refer to ‘z-power’ as the sum of the difference in z-scores calculated using the mean and standard deviation of all metrics (Table S4) between a simulated dataset and its reference dataset(s). The passive and active pooled cases reference each other, and have more statistically different outcomes from the first and second-order analyses, so there is more ‘z-power’ to distinguish a case of live cells between passive and active pooled cases compared to individual active modes. The magnitude of the difference in each metric is figured into the z-power (Table S5). Each mode’s ‘z-power’ is derived from how it differs from the active pooled case and the other two active modes. The total z-power for each case is the sum of the absolute value of z-scores:

$$Z_{power} = \sum_{i=1}^{n} z_i$$

where *i* through *n* are the number of statistically different metrics between the case of interest and its reference case(s). This is used to calculate the MEchanism Index (MEI):

$$MEI = \frac{{Z_{power}}^2}{\sqrt{\sum_{i=1}^{n}(z_i - r_i)^2}} \qquad (Equation\ 16\ from\ main\ text)$$

The MEI is scaled by the Z-power so that the magnitude of the similarity between a test case and the reference is greater for cases with more statistical power to differentiate between them. The Z-power is squared to match the units of the sum-of-squares in the denominator and avoid fractional MEI values. Because the capture case has the lowest z-power, it is less distinguishable from other cases, which is reflected in the MEI calculation. Comparisons between capture cases are statistically powerful, but comparisons between capture MEI values and other mode MEI values are less informative.

#### SUPPLEMENTARY FIGURES AND TABLES

**Supplementary Table 1.** *Comparing matching of base case candidate parameters* to live cell data

| Case | Area KS-value | Area p-value | Sidedness p-value | Sum of p values |
|---|---|---|---|---|
| Base (0.85, 30, 0.003) [Packing, viscosity, CE rate] | 0.07 | 0.41 | 0.18 | 0.59 |
| High viscosity (20) | 0.06 | 0.45 | 0.29 | 0.74 |
| Low viscosity (40) | 0.07 | 0.33 | 0.38 | 0.71 |
| Low packing (0.6) | 0.08 | 0.26 | 0.32 | 0.58 |
| High packing (1.0) | 0.06 | 0.55 | 0.52 | 1.07 |
| Low CE rate (0.002) | 0.09 | 0.22 | 0.49 | 0.71 |
| High CE rate (0.004) | 0.11 | 0.23 | 0.55 | 0.78 |

**Supplementary Table 2.** *Base case parameters for passive case*

| Parameter | Value |
|---|---|
| Viscosity | 0.033 pixels per time step |
| Packing | 1.0 |
| Border speed | 0.003 pixels/time step |
| Diameter | 47.7 pixels |
| Rest range | [0.6, 1.4] |

**Supplementary Table 3.** *Base case parameters for active cases*

| Case | Parameter | Value |
|---|---|---|
| All | Attraction strength | 0.15 |
| Contraction | Angle range | 25 deg from x-axis |
| | Attraction multiplier | 6 |
| Crawling | Speed | 4 pixels/second |
| | Stopping range | 50 pixels from midline |
| Capture | Attraction strength | 8 |
| | Pulling region | 100 pixels around midline |

**Supplementary Table 4.** *Mean and standard deviations for all metrics for all simulated cases*

| METRICS | PASSIVE | | ACTIVE POOLED | | CONTRAC | | CRAWL | | CAP | |
|---|---|---|---|---|---|---|---|---|---|---|
| $z$ | $\bar{x}$ | $\sigma$ | $\bar{x}$ | $\sigma$ | $\bar{x}$ | $\sigma$ | $\bar{x}$ | $\sigma$ | $\bar{x}$ | $\sigma$ |
| Cell ML | -0.014 | 0.046 | -0.071 | 0.061 | -0.044 | 0.062 | -0.085 | 0.055 | -0.08 | 0.058 |
| Cell AP | -0.0034 | 0.055 | -0.075 | 0.059 | -0.053 | 0.064 | -0.083 | 0.053 | -0.082 | 0.055 |
| Cor ML | -0.041 | 0.06 | -0.11 | 0.082 | -0.072 | 0.076 | -0.13 | 0.071 | -0.12 | 0.076 |
| Cor AP | 0.042 | 0.08 | -0.012 | 0.099 | -0.0091 | 0.087 | -0.0087 | 0.1 | -0.015 | 0.11 |
| Cell Area | -0.022 | 0.071 | -0.15 | 0.099 | -0.1 | 0.1 | -0.17 | 0.087 | -0.17 | 0.092 |
| Cor Area | -0.021 | 0.031 | -0.16 | 0.054 | -0.11 | 0.047 | -0.18 | 0.037 | -0.18 | 0.044 |
| T1 freq | 0.279 | 0.183 | 0.23 | 0.144 | 0.2 | 0.18 | 0.21 | 0.13 | 0.17 | 0.098 |
| T1 dwell | 3.2 | 2.5 | 2.6 | 1.8 | 2.4 | 2 | 3 | 2.1 | 2.4 | 1.6 |
| MSD | 0.0075 | 0.0058 | 0.021 | 0.022 | 0.017 | 0.021 | 0.027 | 0.027 | 0.024 | 0.028 |
| ML- | 0.51 | 0.15 | 0.5 | 0.11 | 0.5 | 0.11 | 0.78 | 0.19 | 0.58 | 0.14 |
| ML | 0.46 | 0.17 | 0.47 | 0.14 | 0.47 | 0.14 | 0.8 | 0.17 | 0.61 | 0.17 |
| ML+ | 0.51 | 0.14 | 0.51 | 0.078 | 0.51 | 0.081 | 0.7 | 0.2 | 0.6 | 0.16 |
| AP- | 0.5 | 0.15 | 0.43 | 0.085 | 0.43 | 0.085 | 0.54 | 0.19 | 0.46 | 0.15 |
| AP | 0.47 | 0.2 | 0.49 | 0.19 | 0.49 | 0.19 | 0.57 | 0.23 | 0.53 | 0.19 |
| AP+ | 0.5 | 0.13 | 0.46 | 0.072 | 0.46 | 0.072 | 0.58 | 0.15 | 0.52 | 0.13 |
| Area- | 0.49 | 0.092 | 0.46 | 0.078 | 0.46 | 0.08 | 0.78 | 0.19 | 0.61 | 0.15 |
| Area | 0.49 | 0.17 | 0.5 | 0.22 | 0.49 | 0.22 | 0.8 | 0.16 | 0.64 | 0.15 |
| Area+ | 0.52 | 0.089 | 0.49 | 0.092 | 0.49 | 0.09 | 0.74 | 0.18 | 0.62 | 0.14 |
| PDD ML | 0.48 | 0.18 | 0.41 | 0.15 | 0.42 | 0.16 | 0.4 | 0.15 | 0.41 | 0.15 |
| PDC ML | 0.61 | 0.25 | 0.77 | 0.23 | 0.72 | 0.23 | 0.82 | 0.22 | 0.79 | 0.23 |
| PCD ML | 0.47 | 0.25 | 0.36 | 0.19 | 0.43 | 0.21 | 0.35 | 0.18 | 0.35 | 0.19 |
| PCC ML | 0.62 | 0.2 | 0.67 | 0.19 | 0.65 | 0.21 | 0.69 | 0.19 | 0.7 | 0.18 |
| PDD AP | 0.51 | 0.18 | 0.48 | 0.17 | 0.48 | 0.17 | 0.46 | 0.17 | 0.47 | 0.18 |
| PDC AP | 0.57 | 0.25 | 0.64 | 0.25 | 0.64 | 0.25 | 0.66 | 0.25 | 0.68 | 0.26 |
| PCD AP | 0.51 | 0.25 | 0.48 | 0.25 | 0.5 | 0.24 | 0.49 | 0.24 | 0.47 | 0.25 |
| PCC AP | 0.58 | 0.2 | 0.61 | 0.2 | 0.63 | 0.2 | 0.58 | 0.19 | 0.63 | 0.21 |

**Supplementary Table 5.** *Z-scores and z-power for simulated cases with statistical significance*

| METRICS | PASSIVE | ACTIVE POOLED | CONTRAC | | | CRAWL | | | CAP | | |
|---|---|---|---|---|---|---|---|---|---|---|---|
| *Ref z* | *Ap* | *P* | *Ap* | *Crawl* | *Cap* | *Ap* | *Contrac* | *Cap* | *Ap* | *Contrac* | *Crawl* |
| Cell ML | 0.93 | -1.24 | 0.44 | 0.75 | -0.086 | -0.23 | -0.66 | | | -0.58 | |
| Cell AP | 1.21 | -1.30 | 0.37 | 0.57 | -0.018 | -0.14 | -0.47 | | | -0.45 | |
| Cor ML | 0.84 | -1.15 | 0.46 | 0.82 | -0.132 | | -0.76 | | | -0.63 | |
| Cor AP | 0.55 | -0.68 | | | | | 0.00 | | | | |
| Cell Area | 1.29 | -1.80 | 0.51 | 0.80 | 0.000 | -0.20 | -0.70 | | | | |
| Cor Area | 2.57 | -4.48 | 0.93 | 1.89 | 0.000 | -0.37 | -1.49 | | -0.37 | -1.49 | |
| T1 freq | 0.34 | -0.27 | | -0.08 | 0.408 | | 0.06 | 0.408 | -0.42 | -0.17 | -0.308 |
| T1 dwell | 0.33 | -0.24 | | -0.29 | 0.375 | | 0.30 | 0.375 | | 0.00 | -0.286 |
| MSD | -0.61 | 2.33 | | -0.37 | 0.107 | 0.27 | 0.48 | | | 0.33 | |
| ML- | | | | | | | | | | | |
| ML | | | | | | | | | | | |
| ML+ | | | | | | | | | | | |
| AP- | | | | | | | | | | | |
| AP | | | | | | | | | | | |
| AP+ | | | | | | | | | | | |
| Area- | | | | | | | | | | | |
| Area | | | | | | | | | | | |
| Area+ | 0.33 | | | | | | | | | | |
| PDD ML | | | | 0.13 | -0.067 | | -0.13 | | | -0.06 | |
| PDC ML | -0.70 | 0.64 | -0.22 | -0.45 | 0.130 | | 0.43 | | | 0.30 | |
| PCD ML | | | | | | | | | | | |
| PCC ML | -0.26 | 0.25 | | | | | | | | | |
| PDD AP | | | | | | | | | | | |
| PDC AP | -0.28 | 0.28 | | | | | | | | | |
| PCD AP | | | | | | | | | | | |
| PCC AP | | | | | | | | | | | |
| Z Power | 10.25 | 14.66 | 2.93 | 6.15 | 1.32 | 1.21 | 5.48 | 0.78 | 0.79 | 4.02 | 0.59 |

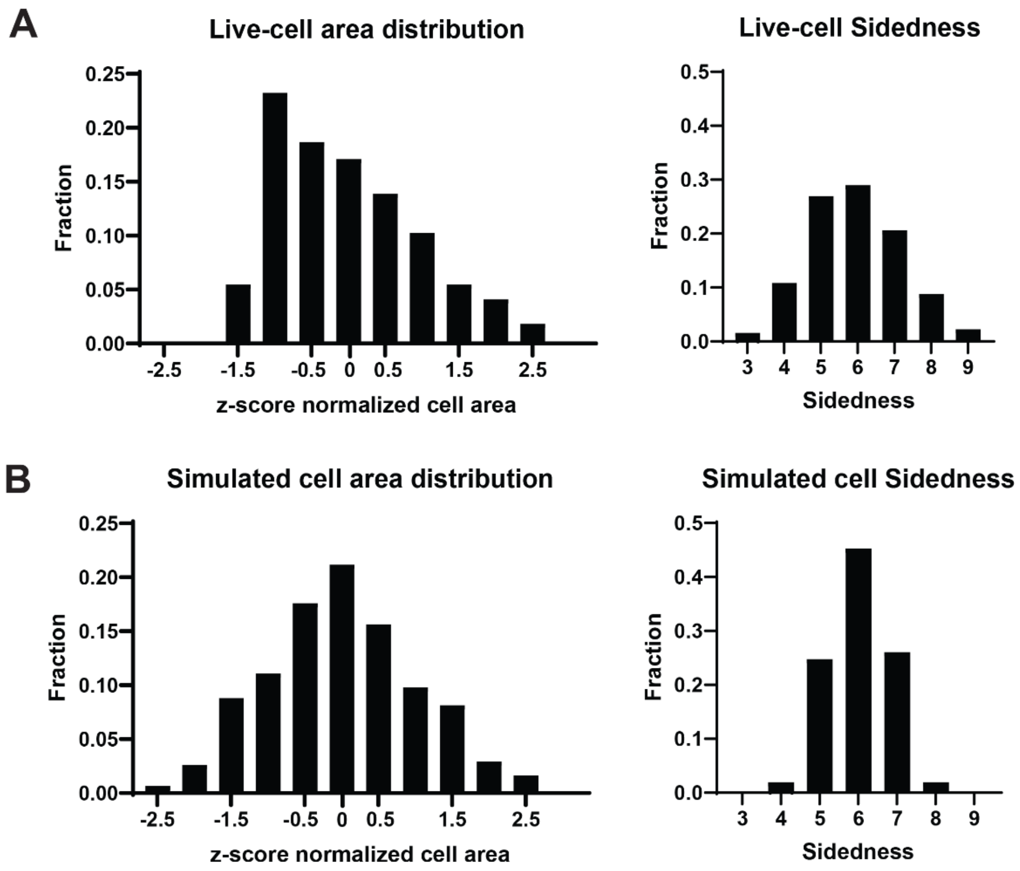


**Supplementary Figure 1.** *Area and sidedness*, supporting Figure 1. [n=1354 cells across 3 explants and 2 whole embryos] (A) Histogram of Z-score normalized distribution of cell area and sidedness in *Xenopus*. (B) Histogram of Z-score normalized distribution of cell area and sidedness in the simulated base case.

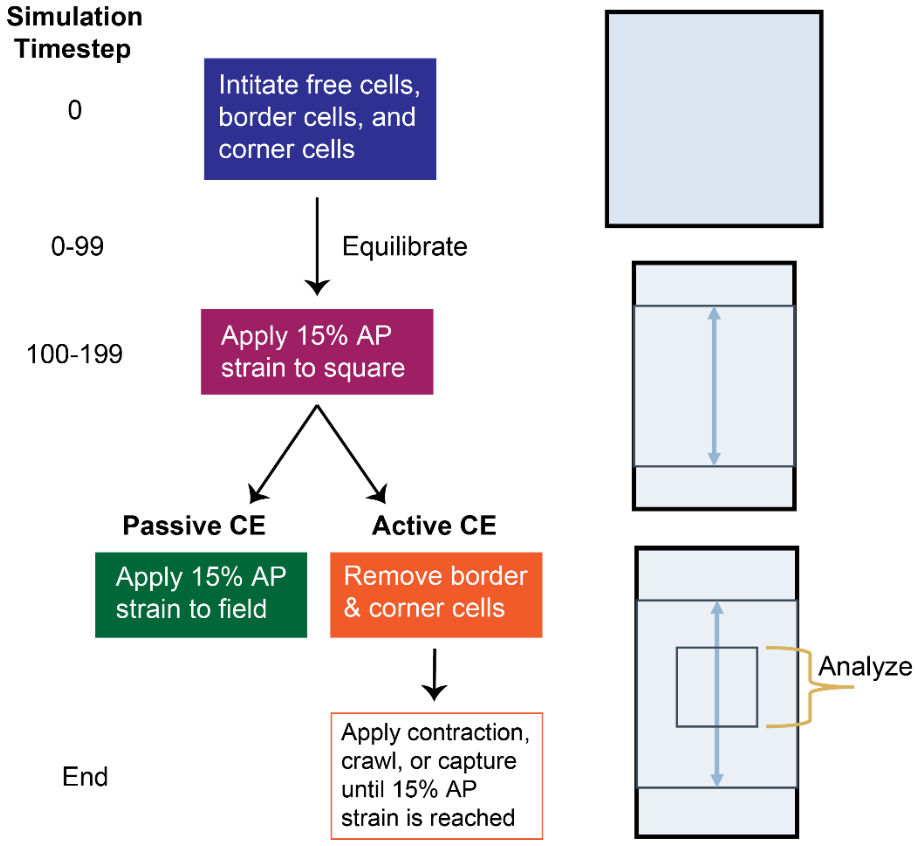


**Supplementary Figure 2.** *Simulation initialization and pre-strain application*, supporting Methods. Steps of simulation initialization and pre-strain to match live-cell tissues that are already undergoing strain at the time of imaging. Active and passive CE-driving forces are separated at the same time step, depending on the case. All simulations are stopped once 15% AP strain is reached in the middle region, and the cells in the middle third are used for all analyses.

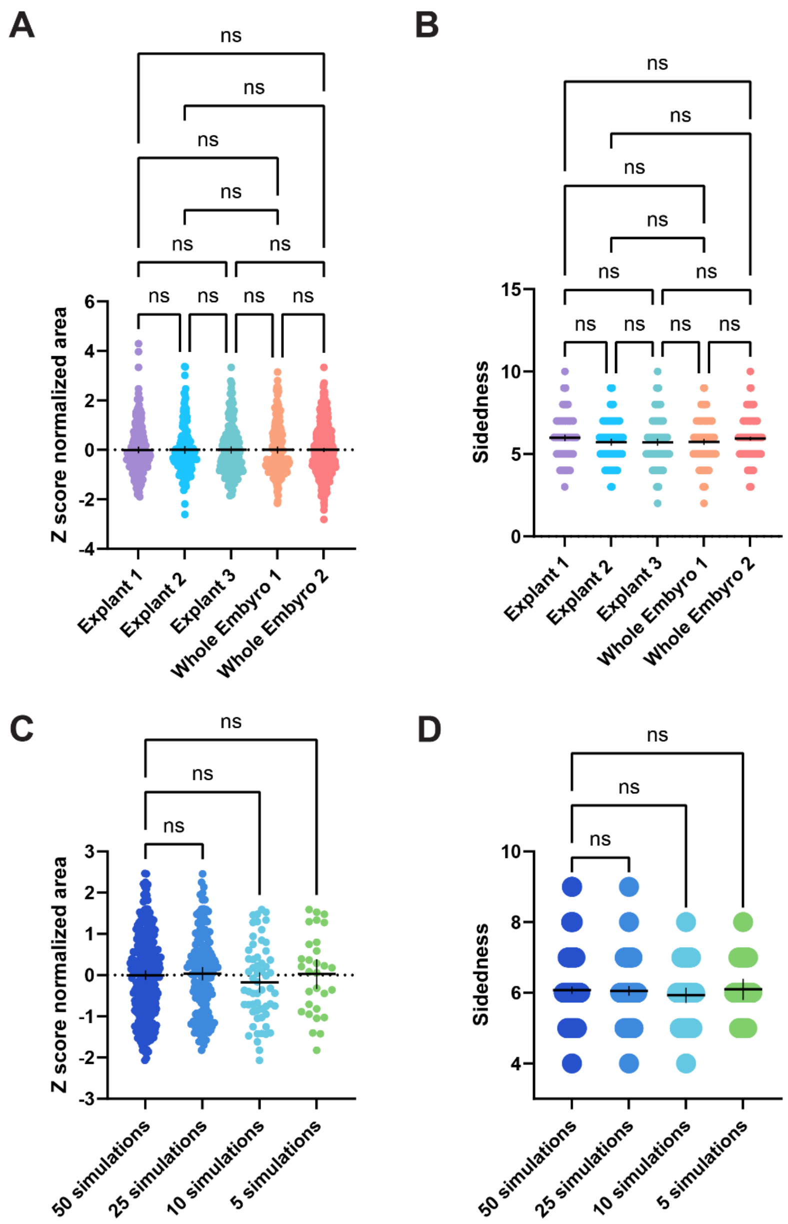


**Supplementary Figure 3.** *Data pooling and sample size selection for live cell data and simulations*, Methods. [n=200 cells (Explant 1), n=177 cells (Explant 2), n=192 cells (Explant 3), n=214 cells (Whole Embryo 1), n=572 cells (Whole Embryo 2); n=294 cells (50 simulations), n=147 cells (25 simulations), n=59 cells (10 simulations), n=30 cells (5 simulations)] (A) Z-score normalized cell areas in *Xenopus*, shown to be not significantly different between individual explants or whole embryo neural plate cells, nor between explants and neural plates. As such, the cells were pooled for analysis. (B) Sidedness of explants and whole embryos showing the same lack of significant differences that allow for pooling. (C) Serial down-sampling of z-score normalized cell areas of the base case of simulations with no differences of areas and sidedness. Ten simulations were chosen to match the area and sidedness of live cells. (D) Serial down-sampling of sidedness of the base case of simulations showing the same trend as areas to support the sample size chosen.

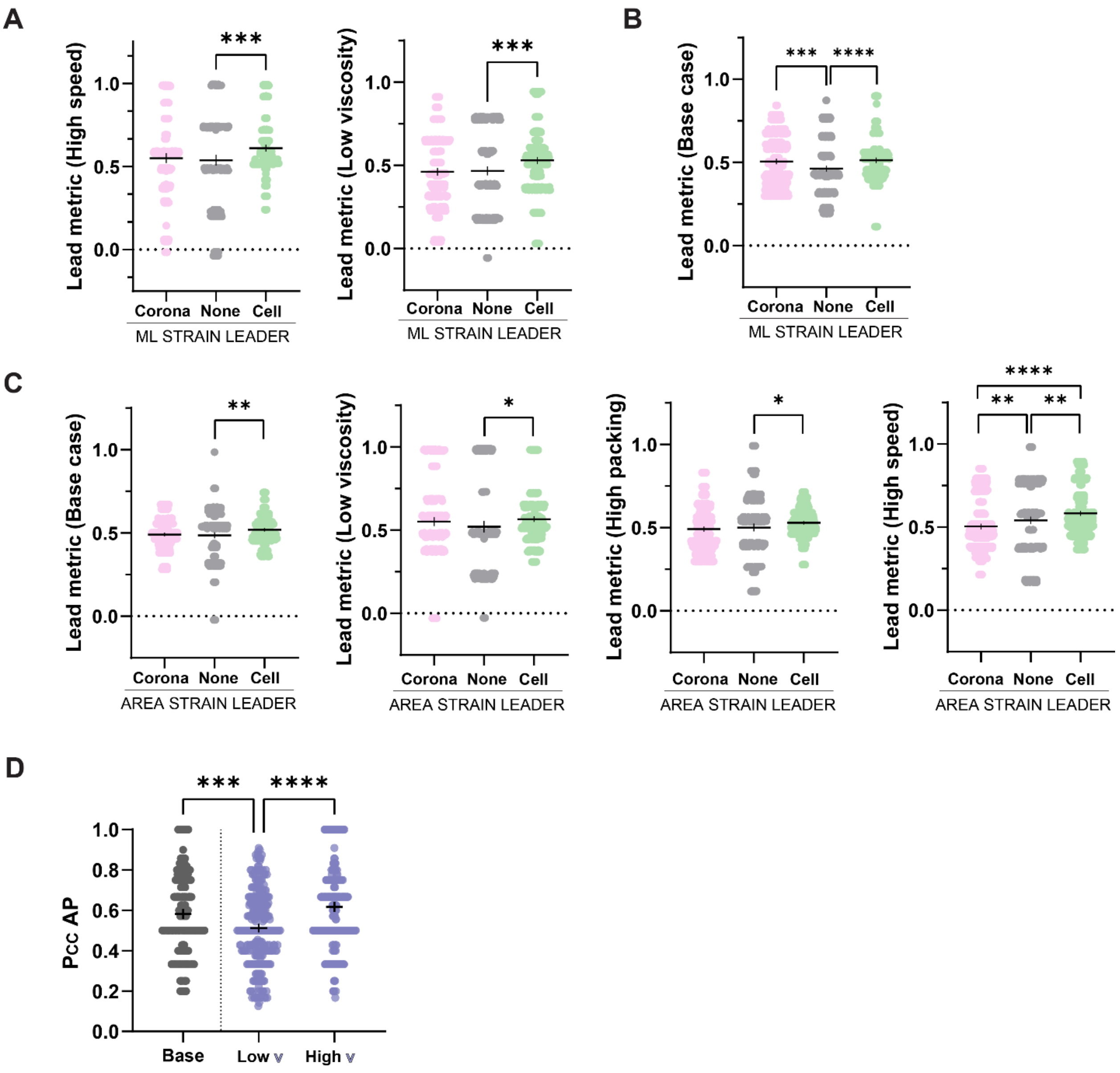


**Supplementary Figure 4.** *Passive case sensitivity,* supporting Figure 3. [sample sizes match Figure 3] (A) Lead metric of high CE speed and low viscosity cases that exhibit cell-led ML strain. (B) The lead metric of the base case showing lack of a strain leader. (C) Base, low viscosity, high packing, and high CE speed cases showing cell-led area strain. (D) Concordance probability in the AP direction for the low and high CE speed cases of the same trend as the ML direction for passive CE cases.

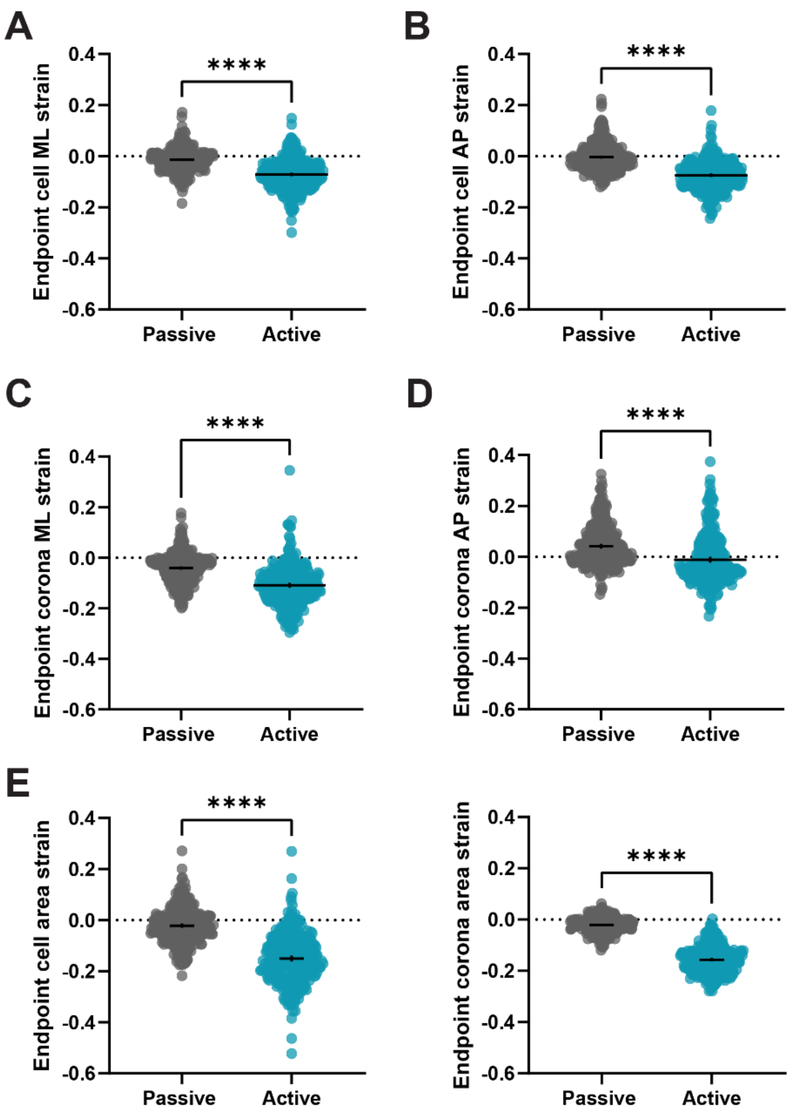


**Supplementary Figure 5.** *Passive case endpoint strains*, supporting Figure 5C. [Sample sizes match Figure 5] (A) Endpoint cell ML strain compared between passive and active pooled cases. (B) Endpoint cell AP strain compared between passive and active pooled cases. (C) Endpoint corona ML strain compared between passive and active pooled cases. (D) Endpoint corona AP strain compared between passive and active pooled cases. (E) Endpoint cell area strain compared between passive and active pooled cases. (F) Endpoint corona area strain compared between passive and active pooled cases.

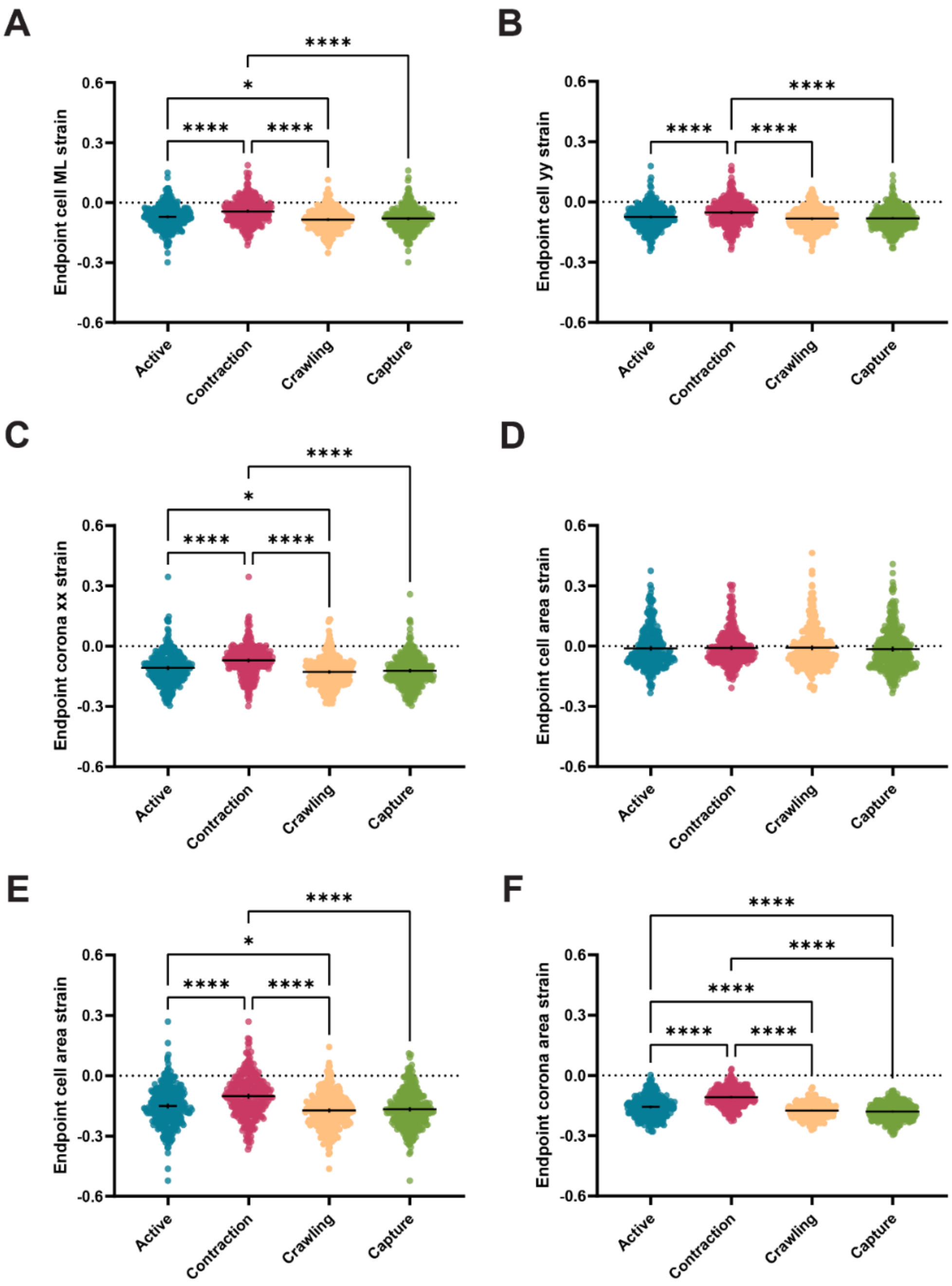


**Supplementary Figure 6.** *Active case endpoint strains*, supporting Figure 6C. [Sample sizes match Figure 6] (A) Endpoint cell ML strain compared between the active pooled case and individual active cases. (B) Endpoint cell AP strain compared between the active pooled case and individual active cases. (C) Endpoint corona ML strain compared between the active pooled case and individual active cases. (D) Endpoint corona AP strain compared between the active pooled case and individual active cases. (E) Endpoint cell area strain compared between the active pooled case and individual active cases. (F) Endpoint corona area strain compared between the active pooled case and individual active cases.

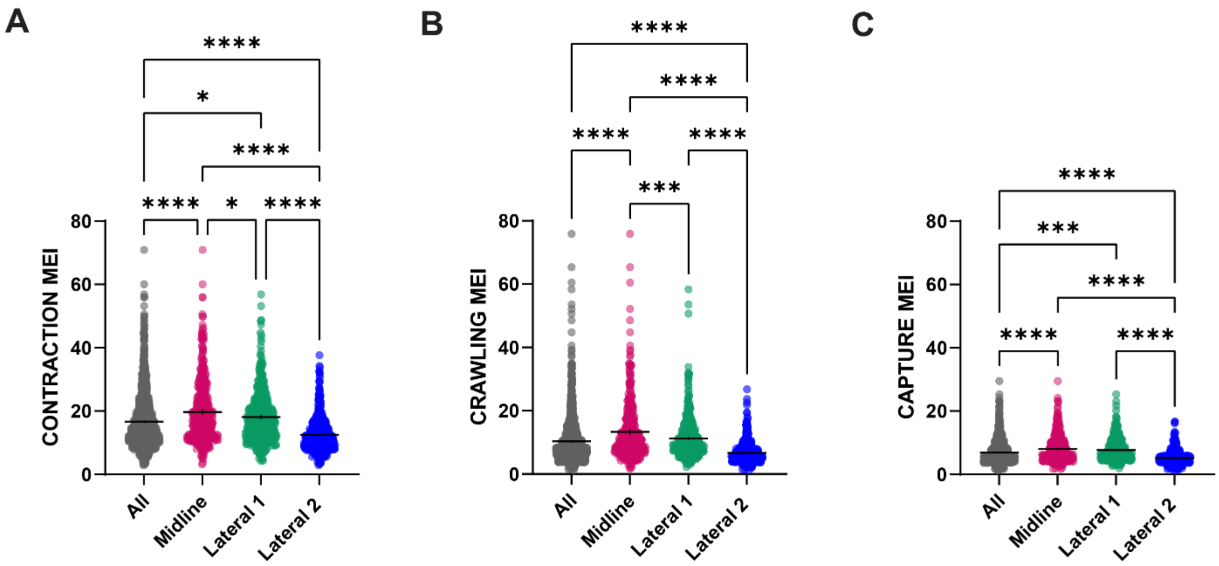


**Supplementary Figure 7.** *MEI for active modes by cell for frog data for all regions*, supporting Figure 7F,H. [Sample sizes match Figure 7] (A) Contraction MEI compared between the frog neural plate as a whole and the midline, lateral 1, and lateral 2 regions. (B) Crawling MEI compared between the frog neural plate as a whole and the midline, lateral 1, and lateral 2 regions. (C) Capture MEI compared between the frog neural plate as a whole and the midline, lateral 1, and lateral 2 regions.

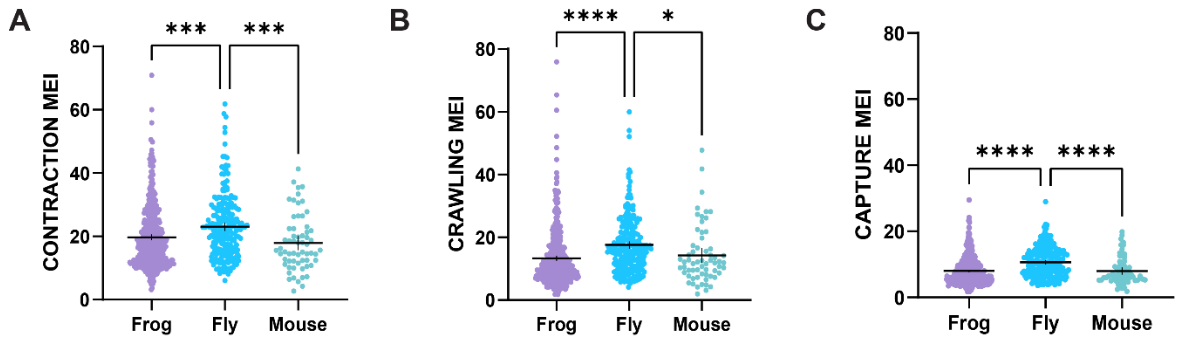


**Supplementary Figure 8.** *MEI for active modes by cell for frog, fly, and mouse data*, supporting Figure 8G. [Sample sizes match Figure 8] (A) Contraction MEI compared between the frog midline, fly, and mouse tissue undergoing CE. (B) Crawling MEI compared between the frog midline, fly, and mouse tissue undergoing CE. (C) Capture MEI compared between the frog midline, fly, and mouse tissue undergoing CE.
